# Genetic architecture of the tomato fruit lipidome; new insights link lipid and volatile compounds

**DOI:** 10.1101/2024.07.08.602461

**Authors:** Anastasiya Kuhalskaya, Xiang Li, Jeongah Lee, Itay Gonda, Julia von Steimker, Mustafa Bulut, Esra Karakas, Josef Fisher, Konrad Krämer, Ruben Vanholme, Leah Rosental, Micha Wijesingha Ahchige, Karolina Garbowicz, Annabella Klemmer, Anne-Kathrin Ruß, Andreas Donath, Alvaro Cuadros-Inostroza, Wout Boerjan, Denise Tieman, Dani Zamir, Harry Klee, Saleh Alseekh

**Author notes:** equal contribution (Anastasiya Kuhalskaya, Xiang Li).

## Abstract

Tomato (*Solanum lycopersicum L*.) fruit flavor is determined by a combination of multiple volatile compounds, including several derived from lipids and fatty acids. Although fruit flavor has been intensively studied, the linkage between lipid metabolism and flavor remains largely undefined. Here, we performed a genome-wide association study (GWAS) and QTL mapping for the fruit lipid content from 550 tomato accessions and 107 backcross inbred lines (BILs) in two consecutive seasons. Over 130 lipid compounds were identified and mapped, allowing for the identification of over 600 metabolic QTL (mQTL). We further described and validated candidate genes associated with lipid content. Among them is a lipase-like protein (TomLLP) whose function was validated *in vivo* using overexpression lines in tomato and knockout mutants in Arabidopsis. We also identified functions for three enzymes: a class III lipase (*Sl-LIP8*), a cyclopropane-fatty-acyl-phospholipid synthase (*CFAPS1*), and Lipoxygenase C (*TomLoxC*). By utilizing knockout lines for *CFAPS1* and CRISPR-Cas9 loss-of-function lines for *Sl-LIP8* and *TomLoxC*, we demonstrated the functional importance of these enzymes in fruit lipid metabolism. Our study provides a comprehensive analysis of the tomato fruit lipidome and insights into key genes that shaped the natural variation in tomato lipid content and their links to flavor-associated volatile compounds.

## Introduction

Tomato *(Solanum lycopersicum L.)* is one of the most economically important crops in the world (Knapp et al., 2004), serving as an important source of micronutrients for the human diet. In recent years, great progress has been made in understanding a wide range of metabolic traits associated with fruit compositional quality (Klee & Tieman, 2018). Lipids in plants have essential structural roles within cells as they are major constituents of membranes and constitute much of the cuticle layer that protects plant outer surfaces (Yeats et al., 2012; Fernandez-Moreno et al., 2017; García-Coronado et al., 2022). In addition, lipids act as signaling molecules. One of the main defense hormones, jasmonic acid, is derived from linolenic acid (Vick & Zimmerman, 1984) and fatty acids have been shown to directly induce the expression of defense-related R genes (Chandra-Shekara et al., 2007). Moreover, lipids serve as precursors for many compounds that contribute to flavor, that regulates attraction, and repulsion of herbivores and ultimately, seed distribution (Wang et al., 2001; Chen et al., 2004; Tieman et al., 2012; Garbowicz et al., 2018).

In addition, so-called fatty-acid-derived volatile organic compounds (FA-VOCs) are important contributors to human liking of food crops (Schwab et al., 2008; Tieman et al., 2017; Cortina et al., 2018). In tomato, several FA-VOCs, including multiple C5, C6, C7, C8, and C10 volatiles, are significantly linked with overall liking and flavor intensity (Chen et al., 2004; J. Zhang et al., 2015; Tieman et al., 2017). Even within the *S. lycopersicum* species there is tremendous variation in FA-VOC content; levels of these chemicals within ripe fruit can vary by several orders of magnitude (Tieman et al., 2017). Knowledge of how these volatiles are synthesized and how the pathways are regulated is important for the development of varieties with superior flavor that do not compromise plant defense.

Genetic mapping has been used to characterize and clone a large number of qualitative and quantitative traits in tomatoes including pathogen resistance (Martin et al., 1993), fruit ripening (Manning et al., 2006), β-carotene formation (Ronen et al., 2000), fruit morphology and size (Frary et al., 2000; J. Liu et al., 2002; Xiao et al., 2008; Rodríguez-Leal et al., 2017). Genetic mapping of metabolite abundances enables the identification of metabolite QTL and potentially provides insights into the complex mechanisms underlying the regulation of metabolic pathways (Chen et al., 2004; Grandillo et al., 2007; H. Li et al., 2013; Wu et al., 2016, 2018; Alseekh et al., 2015; Matsuda et al., 2015; Luo, 2015; Garbowicz et al., 2018; Luzarowska et al., 2020; Brouckaert et al., 2023). Advances in metabolomic profiling coupled with an increase in genomic resources has enabled the identification of numerous mQTLs in tomato, for both primary (Causse, 2004; Overy, 2004; Schauer et al., 2006, 2008; Toubiana et al., 2012, 2015) and secondary metabolites (Rousseaux et al., 2005; Tieman et al., 2006; Minutolo et al., 2013; Rambla et al., 2013, 2016; Alseekh et al., 2015, 2017; Schilmiller et al., 2015; Szymański et al., 2020). However, given the relatively low resolution reached using this approach, cloning of the causal genes can be challenging. Genome-wide association studies (GWAS) provide better QTL resolution (Mitchell-Olds, 2010; Korte & Farlow, 2013). The successful identification of complex traits in various crop species has been achieved by combining linkage QTL mapping using biparental populations and GWAS, which helps overcome the limitations of each approach (Wen et al., 2016; Wu et al., 2016; Garbowicz et al., 2018) In tomato, several GWAS have been conducted, leading to, among other insights, an understanding of the history of tomato breeding and domestication. Examples include the 100-fold increased size of the modern tomato relative to its ancestor (Lin et al., 2014) as well as identification of QTL controlling morphological traits (Shirasawa et al., 2013), volatile compounds (J. Zhang et al., 2015; Tieman et al., 2017), and fruit metabolites (Sauvage et al., 2014; Zhu et al., 2018; X. Li et al., 2020). Nevertheless, utilization of the above approaches to investigate natural variation in the tomato fruit lipidome has not been thoroughly investigated.

Here, we describe large-scale lipid profiling of fruit pericarp tissue extracts of a GWAS panel as well as a *S. neorickii* backcross inbred line (BIL) population. We identified 436 and 175 mQTL using GWAS and linkage mapping, respectively. We identified 384 candidate genes associated with lipid content in fruit. To provide deeper insights into lipid metabolism in tomato fruit and the relationship to volatile compounds, we selected five genes for characterization at the molecular level, namely: acetyl-coenzyme A synthetase (*Solyc06g008920*), *Sl-LIP8* (*Solyc09g091050*) encoding a class III lipase, *CFAPS1* (*Solyc09g090510*) encoding a putative cyclopropane-fatty-acyl-phospholipid synthase, a lipase-like protein *(TomLLP, Solyc03g119980)*, and *TomLoxC* (*Solyc01g006450*), encoding lipoxygenase C. To do so, we created and analyzed CRISPR-Cas9 knockout lines for *CFAPS1*, *Sl-LIP8,* and *TomLoxC,* and overexpression lines of *TomLLP*. In addition, we analyzed mutant lines of the *TomLoxC* Arabidopsis orthologue *(CSE, At1g52760)*. We further performed a correlation-based network analysis of lipids, volatiles, and RNA-Seq expression data across 340 tomato accessions. We were able to identify an additional 85 potential lipid-related genes and provide new insights regarding tomato fruit metabolism.

## Results

### A genome-wide lipidomic profile for tomato fruit

To evaluate the genetic underpinnings of the tomato fruit lipidome, we performed family-based QTL mapping using 107 BILs in addition to GWAS on 550 tomato accessions grown in two harvest seasons (see Supplemental Fig. S1 and Supplemental Data Sets S1-6). Using high throughput UHPLC-MS, we were able to detect and quantify 138 lipid compounds (Fig. 1A, Supplemental Data Set S6), classified into six major classes: seven diacylglycerols (DAGs), 19 digalactosyldiacylglycerols (DGDGs), 15 monogalactosyldiacylglycerols (MGDGs), 24 phosphatidylcholines (PCs), 14 phosphatidylethanolamines (PEs), and 59 triacylglycerols (TAGs). We also quantified 2179 distinct lipophilic compounds from two populations (see Supplemental Data Sets S1-4). For the GWAS panel, a principal component analysis (PCA) of all annotated lipid compounds revealed two main groups, consistent with the evolutionary and domestication relationship; tomato wild species including *S. pimpinellifolium*, and domesticated red-fruited accessions largely overlapped with *S. lycopersicum* var. *cerasiforme* (PCA, Fig. 1B). The abundance of different lipid classes was variable across the three groups (Fig. 1C, Supplemental Fig. S1). The wild tomato species had higher lipid levels compared to cultivated varieties, including glycero-, galacto-, and phospholipids. Moreover, most detected non-annotated lipophilic compounds were more abundant in older tomato varieties compared to domesticated ones (Fig. 1, B-C). In addition to exploring the lipidome profiles in the GWAS panel, we quantified 83 lipids, and 826 distinct lipophilic compounds in fruits from a 107-member *S. neorickii* BIL (Supplemental Data Sets S3 and S4). As expected, the lipid profiles exhibited a large variance across the *S. neorickii* lines (Supplemental Fig. S4).

**Figure 1.**
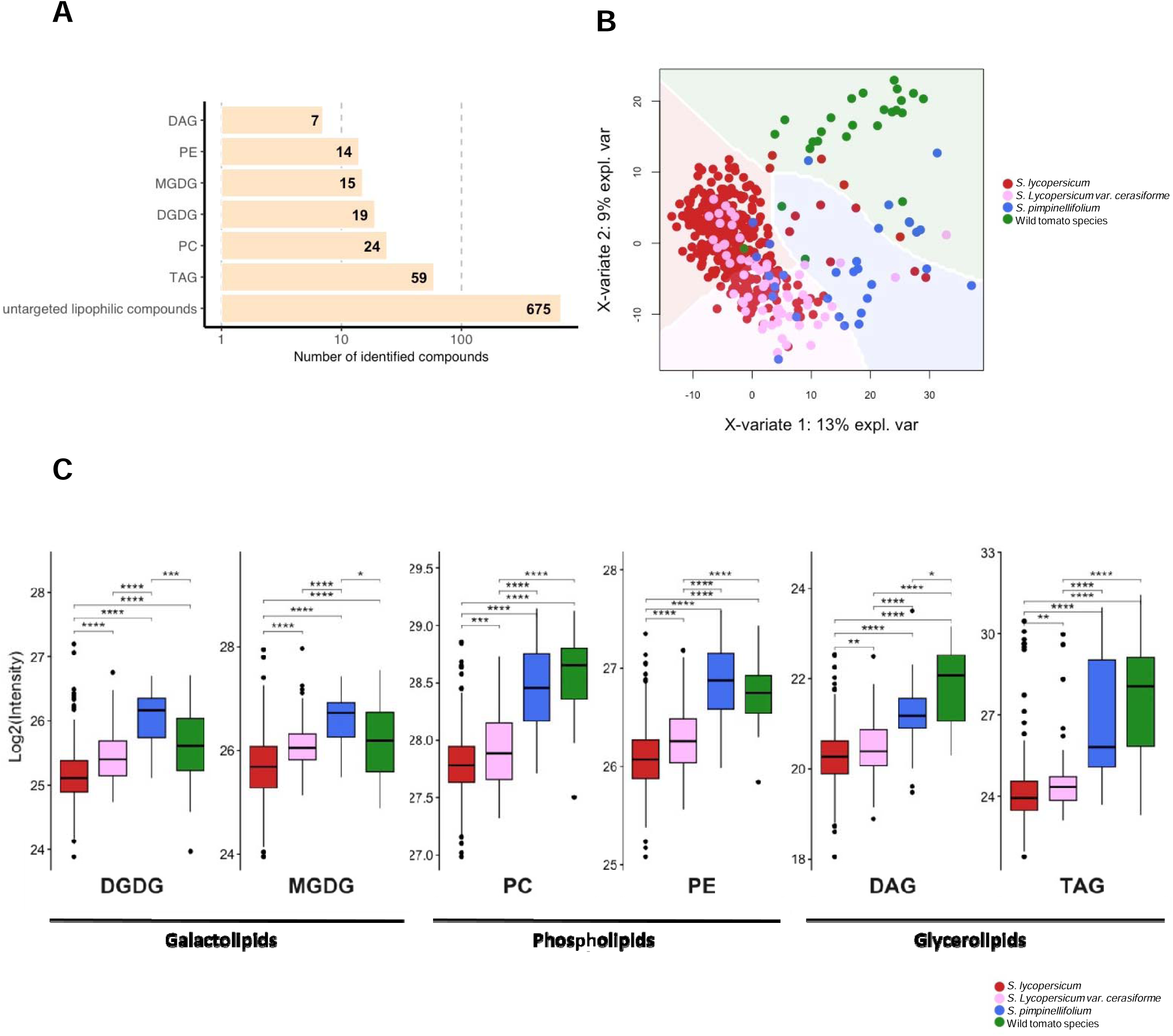
Characterization of Natural Variation in Lipophilic Metabolites Across 550 Different Tomato Accessions. **A)** Numbers of lipid compounds measured by LC-MS in 550 tomato accessions and their compound classes. **B)** PCA of lipid content for tomato lines representing green-fruited wild species (green dots), cultivated varieties (red dots), cherry tomato varieties *S. lycopersicum* var. *cerasiforme* (pink dots), and red-fruited wild accessions of *S. pimpinellifolium* (blue dots). Each dot represents a single accession. **C)** Box plots indicating the average value of all compounds for each lipid class in diverse wild accessions (n = 29), *S. pimpinellifolium* (n = 30), *S. lycopersicum* var. *cerasiforme* (n = 62), and *S. lycopersicum* (n = 398). Significances are indicated by * < 0.05, ** < 0.01, *** < 0.001 using Student’s t-test.

### Genetic foundation of the tomato fruit lipidome

In order to uncover the genetic components of lipid abundances in fruit, GWAS was conducted in two consecutive seasons. We mapped the abundance of 134 annotated and 675 unidentified lipophilic compounds using 1.8 million SNPs (Tieman et al., 2017). In addition, we used 16,526 SNPs generated by genotype by sequencing (GBS) analysis (Zemach et al., 2023) on the GWAS population. We performed QTL mapping on 83 annotated and 826 unidentified lipophilic compounds from BILs using the 10K SolCAP single nucleotide polymorphism chip for linkage mapping.

The GWAS identified 436 significant mQTL (p ≤ 1.0E-04; Supplemental Data Set S6). Linkage mapping using the *S. neorickii* BILs identified an additional 175 significant mQTL in homozygous and heterozygous lines (p ≤ 0.05; Supplemental Data Set S6). Visualizing the distribution of the mQTL in both (GWAS and BILs) populations pointed to a few hotspots for the regulation of a large number of lipids and lipophilic compounds in the tomato genome (Fig. 2; Supplemental Fig. S5), particularly on chromosomes 1, 2, 3, and 6; these represent 12.8%, 15.5%, 11.4%, and 17.3% of the detected loci, respectively (Fig. 2, Supplemental Data Set S6). The vast majority of detected mQTL are for non-annotated lipophilic compounds followed by glycero- and galactolipids. Taken together, we were able to identify 34 conserved lipid mQTL across the tomato genome combining both GWAS and linkage mapping approaches.

**Figure 2.**
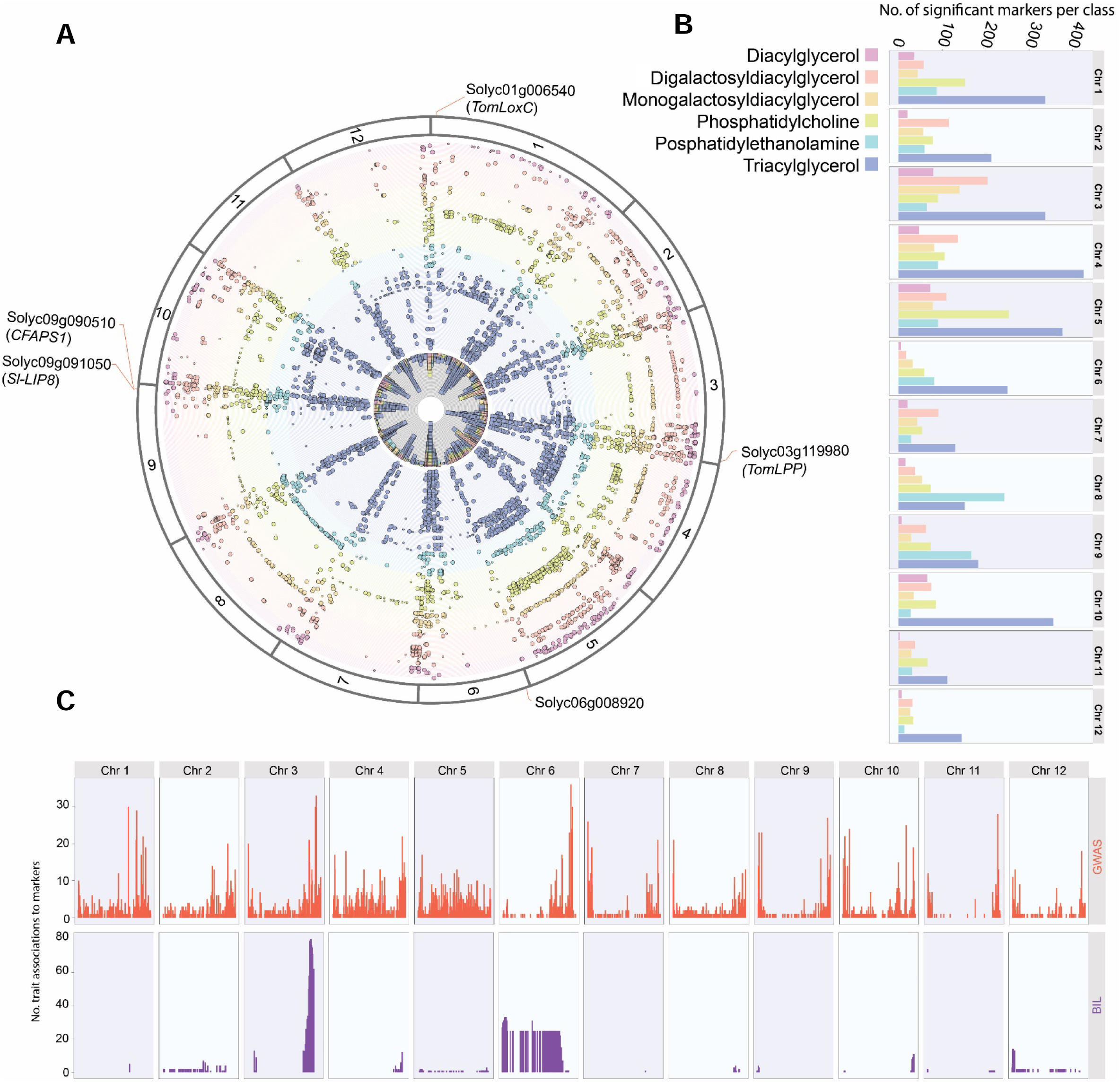
Pleiotropic Map Summarizing Quantitative Fruit Mapping. **A)** Chromosomal distribution of the QTL derived from GWAS represents the combined results from the 2014 and 2015 seasons using SNPs markers generated from Genotype by Sequencing (GBS) and Whole Genome Sequencing (WGS). Colors indicate different lipid classes. The inner circle specifies the amount of lipids mapped to the identified region. QTL harboring candidate genes are highlighted. **B)** Bar charts show the number of significant SNPs associated with each lipid class chromosome-wise. **C)** Number of traits associated with significant markers for GWAS on each chromosome (upper panel) and BIL (lower panel). The corresponding lipid compounds and number of QTL are provided in Supplemental Data Sets S1-5.

### Key genes controlling lipid metabolism in tomato fruit

Having identified mQTL associated with lipid composition in fruit, we investigated several potentially causative genes. Among the identified loci, 384 candidate genes were predicted, based on sequence homology to genes that participate in fatty acid and lipid metabolism (Supplemental Data Set S6). For example, phospholipase D (*Solyc04g082000*), is a potential causal gene for the observed changes in the level of phospho-, glycero- and galactolipids in the QTL on chromosome 4 detected by GWAS in two consecutive seasons. Furthermore, an mQTL located on chromosome 3 identified through both GWAS and BIL mapping, contains a gene annotated as a class III TAG lipase, *Solyc03g123750 (SlLIP2)*. Another gene, *Solyc06g008920,* mapped only in the BIL population (Supplemental Fig. S6), affecting a wide range of long-chain saturated and unsaturated fatty acids, encodes an acyl-CoA synthetase/AMP acid ligase II.

A mQTL detected on chromosome 9 (Fig. 3A) contains an interval of about 50 genes, including *Sl-LIP8* (*Solyc09g091050*), which regulates the biosynthesis of short-chain FA-VOCs by cleaving 18:2 and 18:3 acyl groups from glycerolipids (X. Li et al., 2020) Next, based on the lead SNP, accessions representing the allelic variation was identified for each of the mapped lipids (Fig. 3B). The level of these lipids was plotted according to their domestication status: from *S. pimpinellifolium,* considered to be the ancestor of the modern tomato, *S*. *lycopersicum* var*. cerasiforme*, to the domesticated tomato *S. lycopersicum*, as well as other wild tomato species (Fig. 3C). Of note, the levels of lipids, particularly phosphatidylethanolamine, in *S. pimpinellifolium* differ from the cultivated tomato group. Expression analysis using RNA-Seq data from fruits of the entire GWAS population previously reported (Zhu et al., 2018) showed that *Sl-LIP8* is highly expressed in wild species compared to *S. lycopersicum* var. *cerasiforme* and cultivated tomato (Fig. 3D).

**Figure 3.**
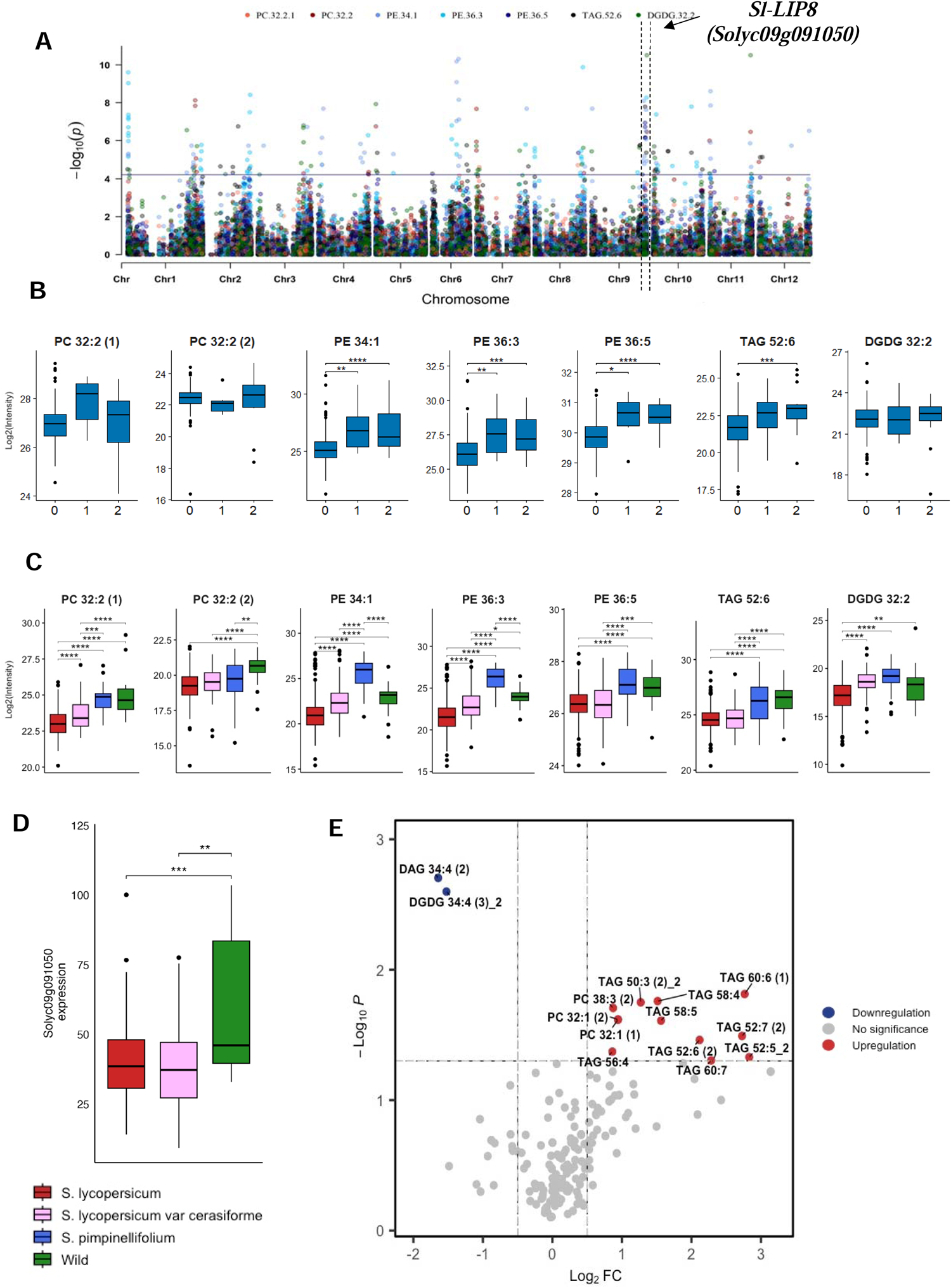
Lipid Contents are Associated with the Locus Harboring *SI-LIP8* (*Solyc09g091050).* **A)** Manhattan plots of the mGWAS results using GBS SNPs data. **B)** Accessions were separated by the lead SNP and the average lipid level was determined. Zero represents the homozygous genotype for the first allele, one represents the heterozygote, and two represents the homozygous genotype for the other allele. **C)** The average lipid level in each of the following groups: *S. lycopersicum* (n = 398)*, S. lycopersicum* var. *cerasiforme* (n = 62), *S. pimpinellifolium* (n = 30), and diverse wild tomato species (n = 27). **D)** *SI-LIP8* transcript levels in fruits of *S. lycopersicum* (n = 258)*, S. lycopersicum* var. *cerasiforme* (n = 56), and diverse wild tomato species (n = 6). **E)** Volcano plot showing the abundance of selected lipids in *SI-LIP8* KO and wild type (Fla. 8059). Lipid levels were calculated as a log2 fold change of Fla. 8059. Significances are indicated by * < 0.05, ** < 0.01, *** < 0.001 using Student’s t-test.

In order to investigate the role of Sl-LIP8 in the tomato fruit metabolome, we generated *Sl-LIP8* CRISPR-Cas9 knockout (KO) lines in the Fla. 8059 background and characterized their metabolites. LC-MS analysis of lipids in fully ripe fruit from KO lines and Fla. 8059 demonstrated notable alterations across various lipid classes (Supplemental Data Set S7). The most substantial changes were observed in the profiles of glycerolipids and phospholipids. Specifically, major differences were observed for polyunsaturated glycerolipids (Fig. 3E) confirming the influence of *Sl-LIP8* on lipid metabolism in tomato fruit.

We also detected another prominent mQTL located on chromosome 9 with a significant impact on the levels of phospho- and glycerolipids using both the GWAS mapping (Fig. 4A) and whole genome sequencing (WGS) SNPs (Fig. 4B). The mQTL harbors two genes encoding putative cyclopropane-fatty-acyl-phospholipid synthases (*Solyc09g090500* and *Solyc09g090510)*. One of these genes, *Solyc09g090510 (CFAPS1),* is expressed in developing fruits (https://bar.utoronto.ca/eplant_tomato/). Accessions responsible for the allelic variation for each of the mapped lipids was identified based on the GBS lead SNP (Fig. 4C) and WGS (Fig. 4D). Lipid levels were plotted along the tomato domestication track, spanning from *S. pimpinellifolium* to *S. lycopersicum* var. *cerasiforme*, and wild tomato species. Notably, the lipid levels, particularly TAG 54:3 and TAG 52:1 across *S. pimpinellifolium* and different wild tomato accessions, differ from those found in the cultivated tomato group (Fig. 4E). Utilizing RNA-Seq data obtained from fruits of the previously characterized GWAS population (Zhu et al., 2018) showed that *CFAPS1 mRNA* was more abundant in cultivated tomatoes (Fig. 4F).

**Figure 4.**
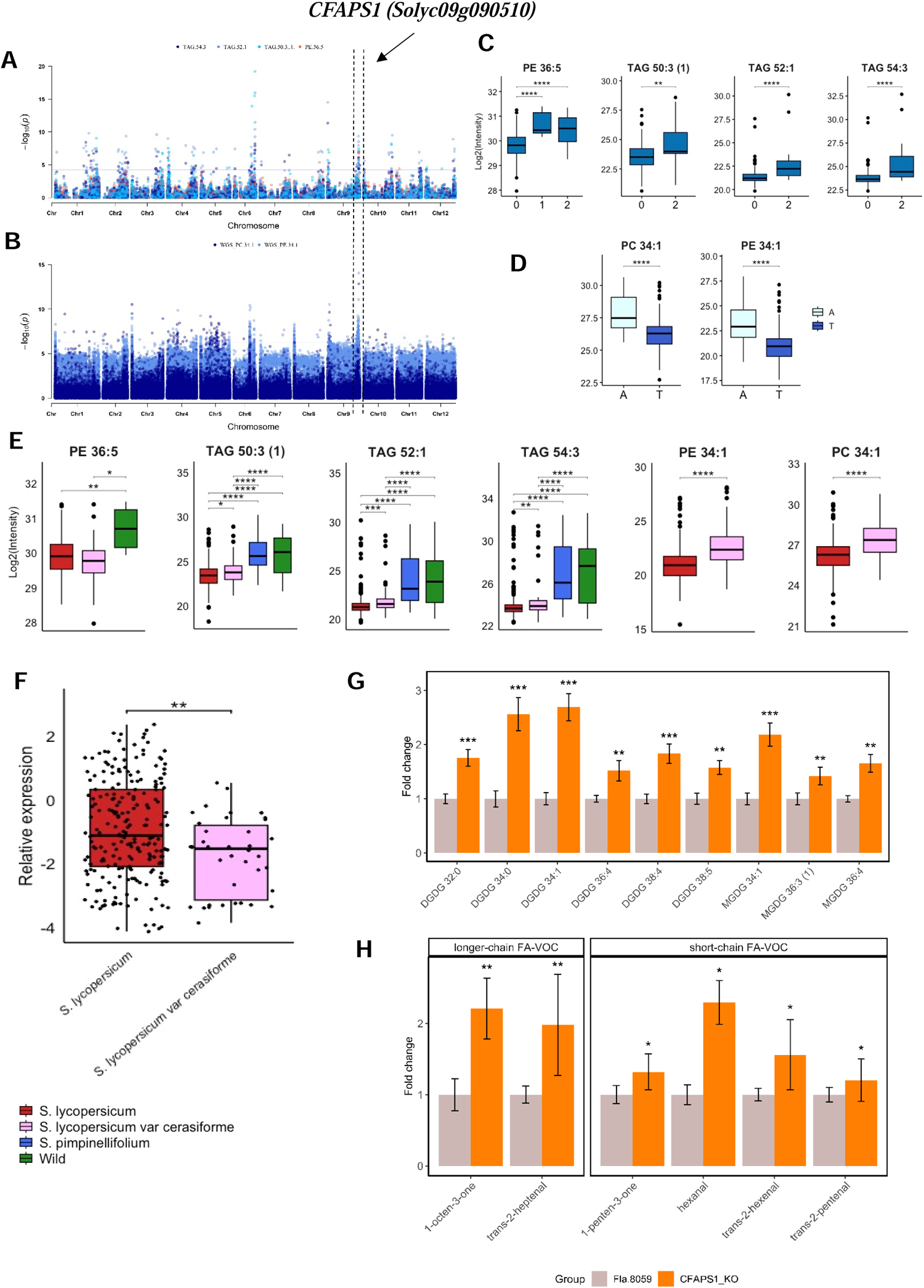
Phospho-, Galacto- and Glycerolipid Contents are Associated with the *CFAPS1 (Solyc09g090510)* Locus. **A)** Manhattan plots of mGWAS using GBS SNPs data. **B)** Manhattan plots of the mGWAS using WGS SNPs data. **C)** Lipid contents in different haplotypes based on the lead mGWAS SNP. Zero is homozygous for the first allele; one is heterozygous; two is homozygous for the second allele. **D)** Lipid analysis of accessions with different haplotypes. **E)** The average lipid level in each of the following: *S. lycopersicum* (n = 398)*, S. lycopersicum* var. *cerasiforme* (n = 62), *S. pimpinellifolium* (n = 30), and diverse wild tomato accessions (n = 27). **F)** *CFAPS1* transcript level in fruits of *S. lycopersicum* (n = 240) and *S. lycopersicum* var. *cerasiforme* (n = 43). **G)** Abundance of selected lipids in *CFAPS1* KO and control (Fla. 8059) fruits. **H)** Abundance of short-chain FA-VOC (C5, C6) and longer-chain FA-VOC (C7, C8) volatiles in *CFAPS1* KO and control (Fla. 8059) fruits. Significances are indicated by * < 0.05, ** < 0.01, *** < 0.001 using Student’s t-test.

We created *CFAPS1* CRISPR-Cas9 KO lines and identified mutants with a deletion of 166 bp and an insertion of 19 bp in the promoter region and coding sequence (exon 1) (Supplemental Fig. S7), which resulted in a premature translation stop at the beginning of the protein. LC-MS analysis of lipidomic profiles in fully ripened fruit from KO and control (Fla. 8059) lines showed significant differences in the contents of various galactolipids (Fig. 4G and Supplemental Data Set S8). In addition, we noticed significant changes in the levels of short-chain (C5, C6) FA-VOCs (Fig. 4H) as well as longer-chain (C7, C8) FA-VOCs (Fig. 4H and Supplemental Data Set S8).

In addition to the above examples, both GWAS and linkage mapping identified a major mQTL influencing the levels of phospholipids and galactolipids at the end of chromosome 3 (Fig. 5, A-E). Linkage disequilibrium analysis of GWAS and recombination breakpoints in a *S. neorickii* BIL population narrowed the mQTL region to 0.33 Mb. This region contains 40 genes, including *TomLLP (Solyc03g119980*), annotated as a lipase-like protein. The population was divided into two haplotypes based on the lead SNP. These two haplotypes exhibited significantly different TAG 50:3 content (Fig. 5C). The *TomLLP* orthologue in Arabidopsis *thaliana*, *CSE* (*At1g52760*), encodes a caffeoyl-shikimate esterase that has been demonstrated to be involved in lignin biosynthesis (Vanholme et al., 2013). CSE has a dual enzymatic activity as both a monoacylglycerol acyltransferase and an acyl hydrolase (W. Gao et al., 2010; Vijayaraj et al., 2012). TomLLP exhibits a strong phylogenetic relationship with caffeoyl shikimate esterase from other plant species (Supplemental Fig. S8 and Supplemental Data Set S9). Next, we selected five *neorickii* BILs covering the QTL interval and measured transcript levels in the same fruit materials used to perform the lipid analysis. We detected higher expression of *TomLLP* in BILs harboring the *S. neorickii* allele compared to the BILs harboring the cultivated allele from the cv. TA209 (Supplemental Fig. S9). To validate our finding, we generated a *TomLLP* overexpression (OE) line in the M82 tomato background carrying the *S. neorickii* allele driven by the figwort mosaic virus 35S promoter (Fig. 5F and Supplemental Data Set S10). Lipid profiling of red ripe fruits from T2 plants showed significant differences in lipid contents between the overexpression line and wild type (Fig. 5G, Supplemental Data Set S11) indicating a role for TomLLP in lipid metabolism and supporting its role as the causative gene associated with the QTL. Furthermore, leaves lipid profiling of three loss-of-function Arabidopsis *CSE* (*At1g52760*) lines demonstrated alterations in the level of multiple lipids belonging to six lipid classes (Fig. 6, A-B, Supplemental Data Set S12). Taken together, the results support a role for TomLLP in tomato fruit lipid metabolism.

**Figure 5.**
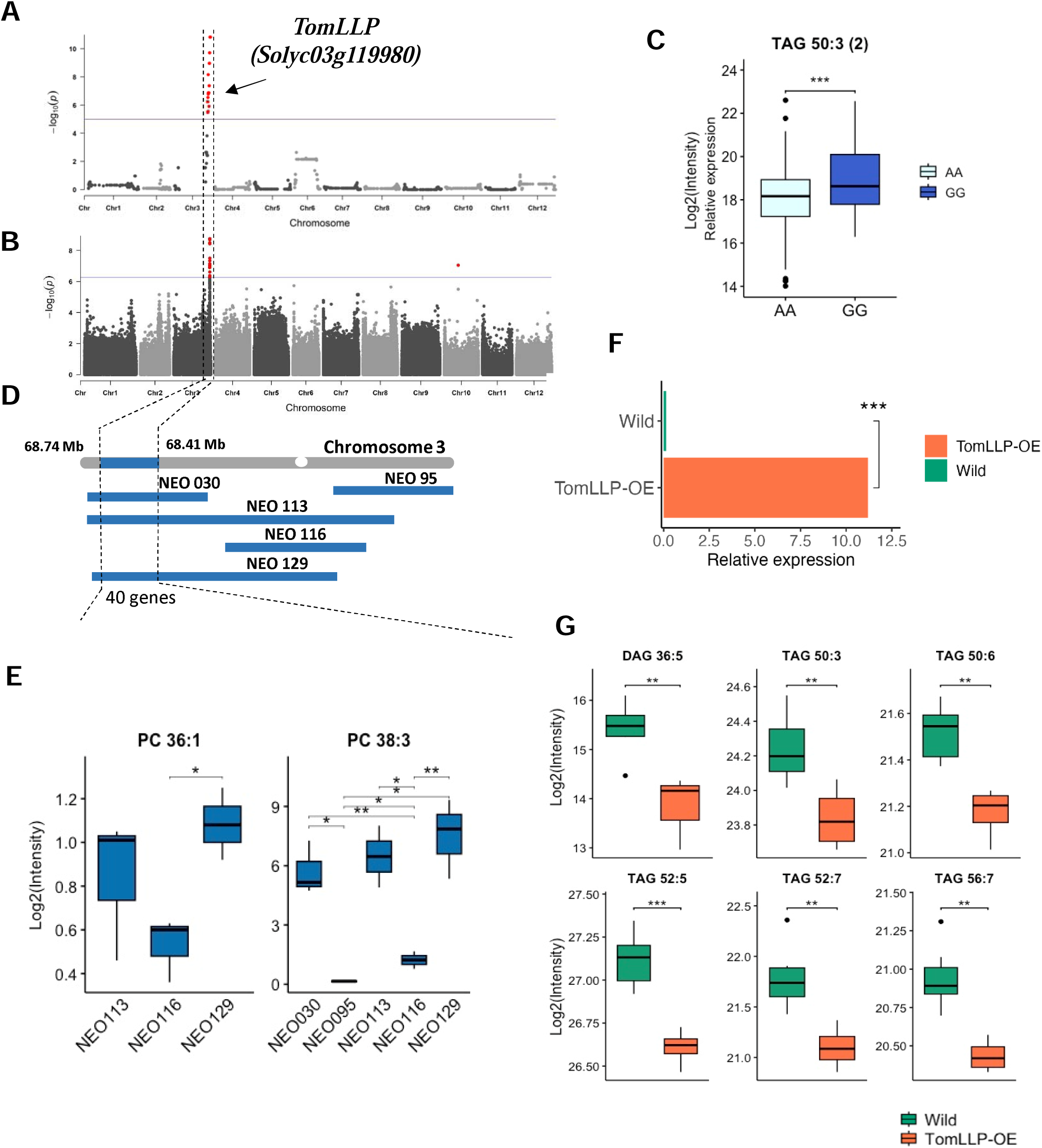
Linkage Mapping Identifies a Role for *TomLLP (Solyc03g119980*) in Fruit Lipid Metabolism. **A)** Association plot of PC 38:3 obtained with linkage mapping using *S. neorickii* BIL population. **B)** Manhattan plot of mGWAS of TAG 50:3 using WGS SNPs data. **C)** Lipid contents of two haplotypes for accessions separated by the lead mGWAS SNP. **D)** *S. neorickii* tomato segments introgressed into cultivated tomato variety TA209 on chromosome 3. **E)** Levels of PC 36:1 and PC 38:3 in BILs sharing the *S. neorickii* introgression on chromosome 3 and BILs with the TA209 background. **F)** *TomLLP* transcript levels in the *TomLLP* overexpression line and wild type M82. **G)** Level of selected lipid in the *TomLLP* overexpression line and M82. Significances are indicated by * < 0.05, ** < 0.01, *** < 0.001 using Student’s t-test.

**Figure 6.**
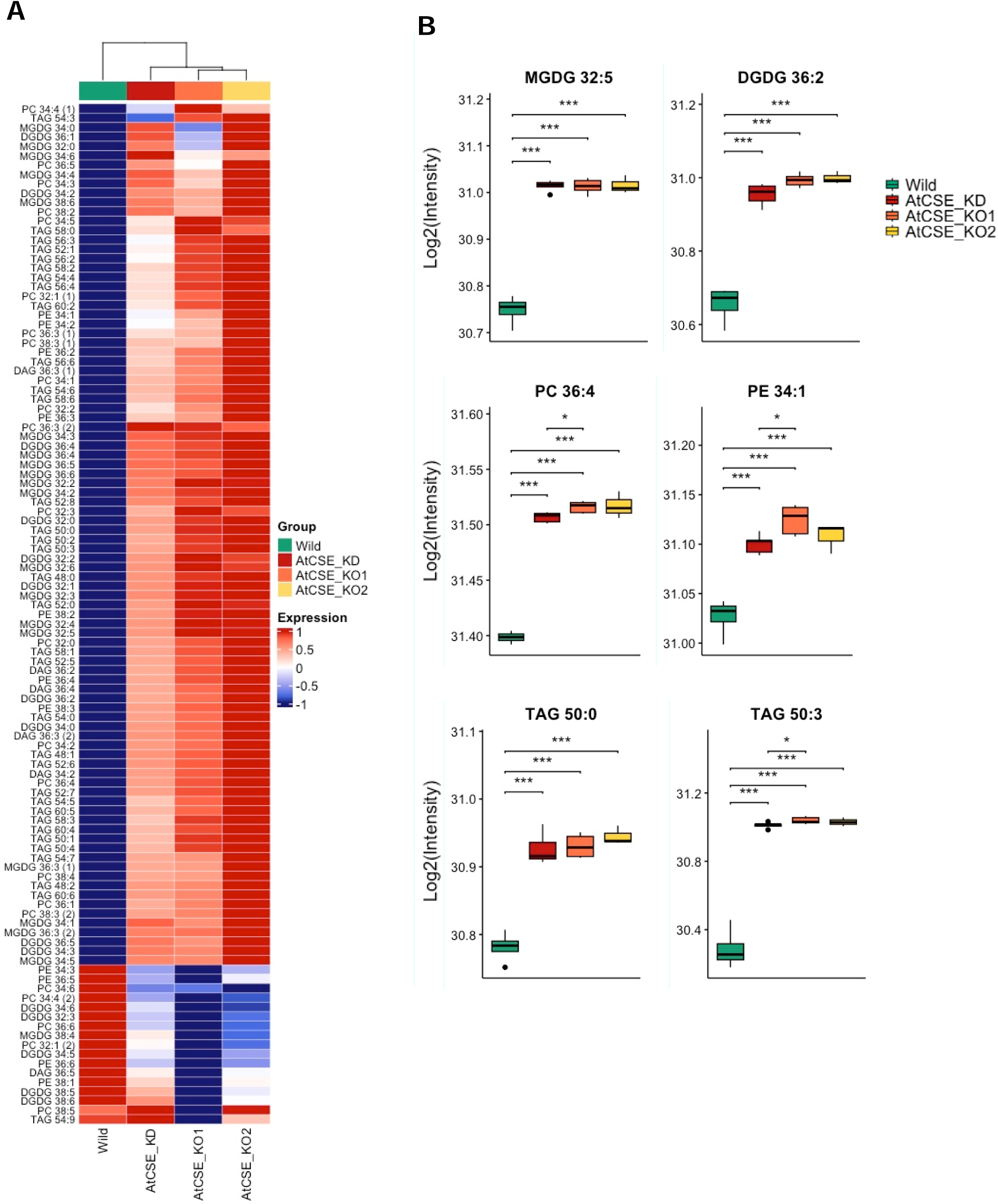
CSE (*At1g52760*) Influences the Lipid Metabolism in Arabidopsis. **A)** Heatmap shows the significant (p ≤ 0.05) changes in lipid levels between wild-type and the *cse* knock-out (KO) and knock-down (KD) lines. **B)** Changes in lipid levels of selected lipids classes between the *cse* KO and KD lines and the wild type.

### Lipoxygenase, a key player affecting tomato fruit lipidome

A robust association between lipids and the locus harboring *TomLoxC* (*Solyc01g006540*) was identified by GWAS (Fig. 7A). Previously TomLoxC was shown to be involved in FA-VOCs production (L. Gao et al., 2019). In order to identify expression differences across the GWAS population, we analyzed previously published *TomLoxC* expression data from 340 accessions (Zhu et al., 2018) and performed eGWAS using 1.8 million SNPs data points (Tieman et al., 2017).A *cis*-eQTL was detected for *TomLoxC* (Fig. 7B) indicating that this gene likely underlies the variation observed in the lipid levels mapped to the *TomLoxC* locus. Accessions responsible for driving the allelic variation for each of the mapped lipids were identified based on the GBS lead SNP (Fig. 7C), and mapped variation in the *TomLoxC* transcript level based on the WGS lead SNP (Fig. 7D). When plotting the median expression level and lipid level associated with the lead SNP we observed that low gene expression coincides with a high level of lipids (Fig. 7, E-F). Interestingly, the levels of some of the lipids mapped to the *TomLoxC* locus were higher in *S. pimpinellifolium* than in cultivated tomatoes (Fig. 7E). These results are consistent with the expression of the mapping panel.

**Figure 7.**
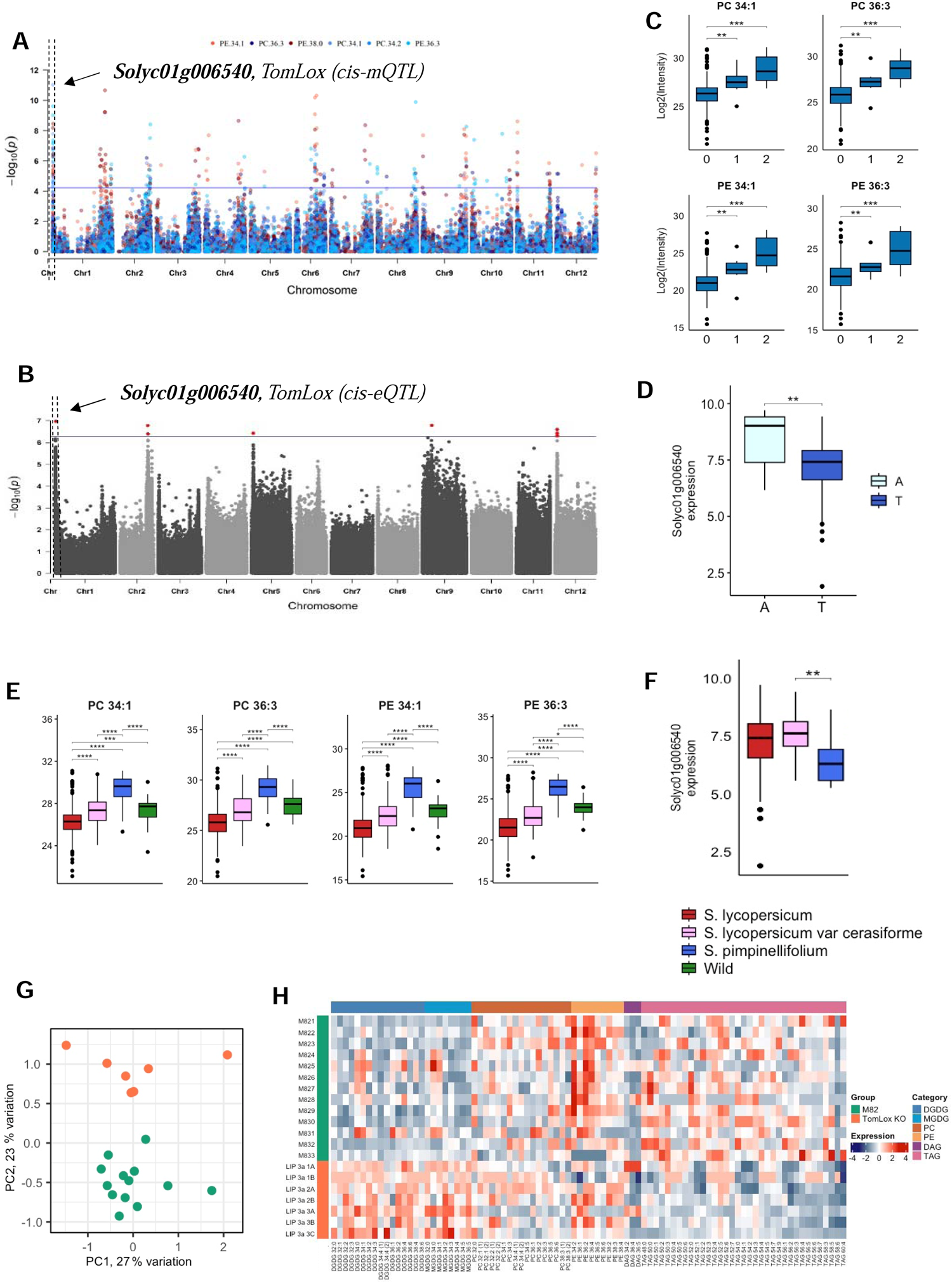
Phospho-, Galacto- and Glycerolipid Content is Associated with the *TomLoxC (Solyc01g006540)* Locus**. A)** Manhattan plots for mGWAS using GBS SNPs data. **B)** Manhattan plot for eGWAS using WGS SNPs data. **C)** Lipid contents for a group of accessions separated by the lead mGWAS SNP. Zero, homozygous for the first allele; one, heterozygous; two, homozygous for the second allele. **D)** Lipid contents for accessions grouped by the lead eGWAS SNP. **E)** Average lipid levels for *S. lycopersicum* (n = 398)*, S. lycopersicum* var. *cerasiforme* (n = 62), *S. pimpinellifolium* (n = 30), and diverse wild tomato species (n = 27). **F)** *TomLoxC* transcript level in fruits of *S. lycopersicum* (n = 258)*, S. lycopersicum* var. *cerasiforme* (n = 56), *S. pimpinellifolium* (n = 6). **G)** PCA plot of lipid levels in *TomLoxC* KO and wild type M82. **H)** Heatmap representing the abundance of short-chain FA-VOC (C5, C6) and longer-chain FA-VOC (C7, C8) in *TomLoxC* KO and wild type M82. Significances are indicated by * < 0.05, ** < 0.01, *** < 0.001 using Student’s t-test.

To provide further insight into the role of *TomLoxC* in lipid metabolism and biologically validate our results, we generated *TomLoxC* CRISPR-Cas9 KO lines in the M82 background. LC-MS lipidomic profiling of red ripe fruit from KO lines and M82 revealed significant changes in a wide range of lipid classes (Fig. 7, G-H, Supplemental Data Set S13). The largest changes were observed in galactolipids followed by phospho- and glycerolipids. Specifically, large differences were observed for polyunsaturated galactolipids (e.g. DGDG 32:2, DGDG 32:3, DGDG 34:3; DGDG 36:6, MGDG 34:2, MDGD 34:5, MGDG 36:5) (Fig. 7H). These observations provide further support for an important role for *TomLoxC* on fruit lipid abundance.

### Transcript-metabolite correlation-based network

In recent years, large-scale gene expression and metabolite data sets have been published for tomato fruit. We used the RNA-Seq data for 340 tomato accessions (Zhu et al., 2018), volatile profiling for 398 tomato accessions (Tieman et al., 2017), and the lipidome data on the GWAS panel obtained in both seasons generated in this study. We constructed a transcript-metabolite correlation-based network using these data (Fig. 8). The result showed that the large majority of the overall correlations were positive (97%) (Supplemental Data Set S14). Negative correlations were observed between phospholipids, glycerolipids, and genes involved in lipid degradation processes and fatty acid oxidation. FA-VOCs were correlated with genes taking part in lipid biosynthesis, signaling, and degradation processes. Genes with lipoxygenase activity showed a connection with different classes of polyunsaturated lipids. For example, TomLoxC, involved in FA-VOCs biosynthesis (Chen et al., 2004; Tieman et al., 2017; Klee & Tieman, 2018) was positively correlated with phospholipids PC 36:2 (*r*=0.32), and PE 36:5 (*r*=0.33). Of interest*, TomLLP* showed negative interaction with the FA-VOC hexyl alcohol (*r*=-0.31). Finally, four genes (*Solyc02g071700, Solyc02g077430, Solyc06g053900,* and *Solyc08g063090*) showed positive correlations with phospho-, galacto-, and glycerolipids and these genes fell within the mQTL intervals we identified as candidate genes in this study (Supplemental Data Set S6).

**Figure 8.**
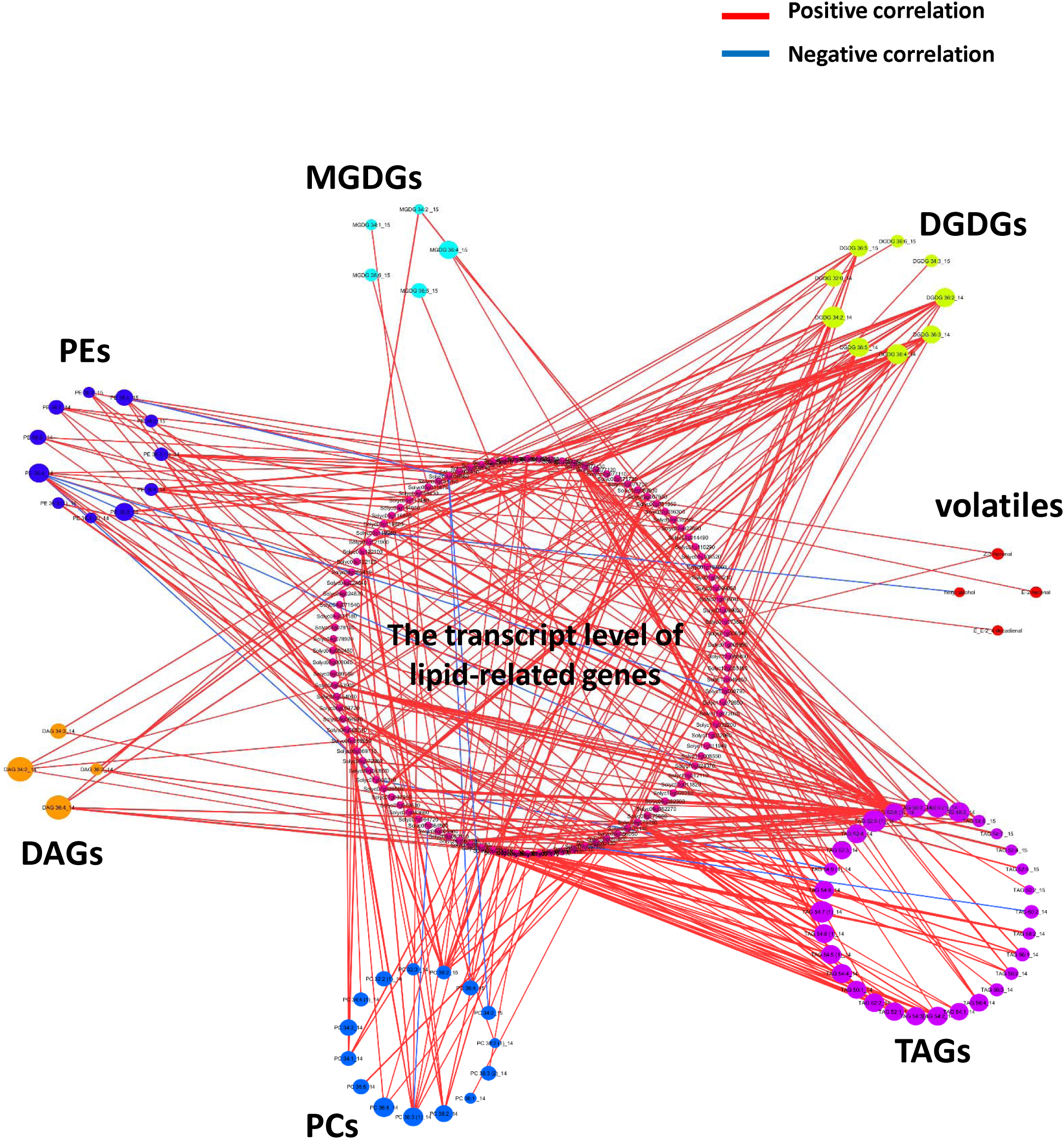
Metabolite-Transcript-Volatile Correlation-based Network. Each node represents a metabolite or gene transcript; edges connecting two nodes show a correlation (R ≤ - 0.3, or R ≥ 0.3) between the two nodes. In total, the network is composed of 185 nodes and about 335 edges assembled into three large groups: lipophilic metabolites comprise 74 nodes, gene expression data have 107 nodes, and four nodes for volatile organic compounds (VOCs; Supplemental Data Set S14). There are 672 genes with homology to genes known to be involved in lipid metabolism Garbowicz et al., 2018). Transcript levels were used to construct the network (Zhu et al., 2018), VOC data (Tieman et al., 2017), and all other lipid metabolites derived from the current study.

## Discussion

In recent years, the application of many metabolic GWAS studies has focused on dissecting the genetic architecture underlying regulation and biosynthesis of metabolic pathways (Alseekh et al., 2021). However, application of this approach to investigate the fruit lipidome of tomato and its link to fruit flavor has not been performed. Here, through a combination of GWAS and linkage analysis, we identified over 600 mQTL and many genes that affect lipid composition in tomato fruit. In order to validate the results, we determined the functions of four lipid biosynthesis genes by transgenic analysis.

Metabolite GWAS has become increasingly common in recent years, particularly for lipids (Fang & Luo, 2019; Luzarowska et al., 2020; Brouckaert et al., 2023). These studies have revealed novel associations between structural genes and lipids. In tomato, limited work based on the natural variation in a bi-parental population has been carried out to study cuticle lipids (Yeats et al., 2012; Fernandez-Moreno et al., 2017). Here, we utilized both *S. neorickii* BILs and a GWAS panel (Supplemental Fig. S1) to map the lipidome of tomato fruit pericarp (Fig. 1), revealing genetic associations between lipids, volatiles, and gene expression (Fig. 2 and Supplemental Fig. S5). We examined the variation in lipid levels among cultivated and wild species (Fig. 1, A-B). Wild tomato accessions had higher lipid levels as compared to cultivated tomatoes (Fig. 1C). Domestication of tomato has resulted in a narrower genetic and phenotypic variation in cultivated species (Ranc et al., 2008; Bergougnoux, 2014; Blanca et al., 2015).The utility of exotic germplasm as a source of new traits has been extensively exploited (Bessey, 1906; Zamir, 2001; McCouch, 2004; Doebley et al., 2006; Fernie et al., 2006; Grandillo et al., 2007). For example, introgression populations containing portions of the genomes of the wild tomato relatives *S. pennellii* and *S. neorickii* into cultivated tomato have provided useful tools to explore multiple genes with roles in morphological and metabolic traits (Schauer et al., 2008; Chitwood et al., 2013; Alseekh et al., 2015; Brog et al., 2019), including lipids (Garbowicz et al., 2018). The large differences in chemical composition between fruits of the different species mean that introgressions of defined segments from those wild relatives frequently result in major perturbations of metabolic pathways. Our integrative approach using two types of populations has been proven to be particularly useful, allowing us to cross-validate results (Fig. 1-2). We identified 436 mQTL and 175 mQTL using GWAS and linkage mapping (*S. neorickii* BILs) approaches, respectively (Supplemental Data Set S6). Thirty-four mQTL and 38 candidate lipid-related genes were common between the two mapping populations.

The mGWAS uncovered an association between the levels of multiple lipid classes and the genomic locus harboring *Sl-LIP8* (as shown in Fig. 3A). Additionally, *Sl-LIP8* mutants showed significantly increased levels of several TAGs (as depicted in Fig. 3E). *Sl-LIP8* encodes a class III lipase, which has been demonstrated to cleave TAGs and DAGs, leading to the subsequent release of volatile compounds (Garbowicz et al., 2018). Moreover, Garbowicz et al. precisely mapped the mQTL containing *Sl-LIP8* to a shared region within *S. pennellii* introgression lines (IL 9-3, IL 9-3-1, and IL 9-3-2), where a noticeable reduction of levels of DAGs, DGDGs, MGDGs, and TAGs were observed in comparison to control (Garbowicz et al., 2018). Sl-LIP8 participates in the biosynthesis of important flavor-associated FA-VOCs (X. Li et al., 2020) (Supplemental Fig. S10B). Therefore, Sl-LIP8 acts as a bridge connecting lipid and volatile metabolism presumably through the cleavage of glycero- and phospholipids, leading to the release of free fatty acids which are further metabolized to short-chain FA-VOCs (Garbowicz et al., 2018; X. Li et al., 2020) (Supplemental Fig. S10B). Previous results indicated that the transcript level of *Sl-LIP8* is higher in the *S. lycopersicum* cv. M82 tomato variety than in *S. pennellii* (Garbowicz et al., 2018), potentially due to the structural variation in the promoter regions (Kuhalskaya et al., 2020). In our study, we observed increased *Sl-LIP8* expression in wild tomato species such as *S. habrochaites, S. arcanum, S. chmielewskii* and *S. peruvianum*. These species are positioned closer to *S. lycopersicum* on the domestication track than *S. pennellii* (Koenig et al., 2013; Bolger et al., 2014; Lin et al., 2014). Additionally, the elevated *Sl-LIP8* expression was correlated with various lipid alterations (Fig. 3, C-D).

The GWAS mapping uncovered mQTL harboring the *CFAPS1* gene, which is responsible for observed changes in the level of various lipid classes and of FA-VOCs (Fig. 4, A-B). *CFAPS1* exhibited higher transcript levels in cultivated tomato accessions with corresponding differences in the lipid levels (Fig. 4, E-F). The *CFAPS1* gene encodes a cyclopropane-fatty-acyl-phospholipid synthase, previously undescribed in tomatoes. In *E. coli*, this enzyme introduces cyclopropane rings into unsaturated membrane phospholipids, resulting in generation of cyclopropane fatty acids (Cronan & Luk, 2022). In plants, these unusual cyclopropane fatty acids can be utilized for TAGs and DAGs synthesis (Shockey et al., 2018), which can then be used as substrates to produce volatile compounds (Garbowicz et al., 2018) (Supplemental Fig. S10, B-C). Our study demonstrated that the tomato *CFAPS1* targets monounsaturated or polyunsaturated TAGs and membrane lipids (Fig. 4). Membrane lipids in plants include phospholipids and galactolipids (Li-Beisson et al., 2013). The *CFAPS1* mutants showed a significant increase in the levels of various galactolipids (Fig. 4G). Furthermore, the KO lines exhibited substantial variations in the composition of short-chain (C5, C6) and longer-chain (C7) FA-VOCs, with remarkable increases of hexanal (C6), *E*-2-hexenal (C6), *E*-2-heptenal (C7), and 1-octene-3-one (C8) (Fig. 4H). In tomato fruit, certain C5, C6, C7, C8, and C10 FA-VOCs, exhibit significant correlations with flavor intensity and overall preference (J. Zhang et al., 2015; Tieman et al., 2017). Thus, our validation establishes *CFAPS1* as the causal gene responsible for the natural variation in lipid abundances associated with this locus. Moreover, our study demonstrates that *CFAPS1* is another bridge connecting lipid and volatile metabolic pathways (Supplemental Fig. S10C).

Another gene that was shown to participate in synthesis of FA-VOCs, through both lipase-dependent and lipase-independent pathways, is *TomLoxC* (Klee, 2010; Klee & Tieman, 2013; Mwenda & Matsui, 2014; Garbowicz et al., 2018; Zhao et al., 2019) (Supplemental Fig. S10C). Quantitative variation in several lipids was mapped to the *TomLoxC* locus (Fig. 7A). Further support for *TomLoxC* as the causal gene is provided by a *cis*-eQTL (Fig. 7B). *TomLoxC* has higher transcript abundance in cultivated varieties (Fig. 7F). Earlier research documented that *TomLoxC* is a chloroplast-targeted lipoxygenase active during ripening of tomato fruit. *TomLoxC* utilizes both linoleic and linolenic acids as substrates, resulting in the production of flavor-associated short-chain FA-VOCs (Chen et al., 2004; Shen et al., 2014). Lipid profiling of a *TomLoxC* loss-of-function line revealed that *TomLoxC* is associated with significant increases in multiple lipid species with the most significant changes occuring in galactoplipids (Fig. 7H). Thus, the association of various phospholipids with the *TomLoxC* gene and accumulation of galacto-, phospho-, and glycerolipids in the KO lines strongly suggests a role for this enzyme in chloroplast lipid degradation during fruit ripening concomitant with the release of free FAs that are the precursors for ripening-associated FA-VOCs (Supplemental Fig. S10D). Taken together, we validated a role for *TomLoxC* as a causal gene responsible for at least part of the natural variation of lipid abundance in tomato and their close relatives.

Our results showed that, *Sl-LIP8, CFAPS1,* and *TomLoxC* provide a connection between lipid metabolism and FA-VOC biosynthesis. Several of these FA-VOCs have been demonstrated to affect consumer preferences (Tieman et al., 2012, 2017; Cortina et al., 2018). The *TomLoxC* locus appears to be under positive selection within a domestication sweep (Lin et al., 2014). Thus, those genes are important targets for breeding tomato with improved flavor.

We also identified a region containing *TomLLP* on chromosome 3 that affects the levels of phospholipids and galactolipids using both mapping populations (Fig. 5, A-B). *TomLLP* is annotated as a lipase-like protein, and its Arabidopsis orthologue (*At1g52760*), caffeoyl-shikimate esterase (CSE) participates in lignin and lipid biosynthesis (Vanholme et al., 2013) (Supplemental Fig. S10E). Additionally, prior research has demonstrated that *At1g52760* exhibits lysophospholipase activity, utilizing lysophospholipids as substrates, which has a role in phospholipid metabolism (W. Gao et al., 2010; Vijayaraj et al., 2012; Miao et al., 2019) and also has acyltransferase activities, facilitating the synthesis of diacyl- and triacylglycerol (Vijayaraj et al., 2012). Phylogenetic analysis revealed that *TomLLP* clusters closely with the caffeoyl shikimate esterases found in *S. tuberosum,* and *Capsicum Annuum*, rather than with other tomato lipases (Supplemental Fig. 8). Overexpressing *TomLLP* in tomatoes resulted in decreased glycerolipid contents (Fig. 5, F-G), while lipid profiling of three Arabidopsis loss-of-function *cse* mutants (*At1g52760)* exhibited a significant accumulation of multiple lipid species that belong to six lipid classes (Fig. 6, Supplemental Data Sets S11-12). Our results illustrate that both *TomLLP* and *CSE* function in lipid metabolism. Further study is needed to explore if *TomLLP* also has a dual function as does *CSE*.

We performed correlation-based network analysis between lipophilic compounds, transcript abundance of lipid-related genes, and levels of FA-VOCs (Fig. 8). Our analysis showed positive interactions between lipophilic compounds and the expression of lipid-related genes in various metabolic pathways. We identified several lipid-related genes correlated with multiple lipid classes and several lipophilic compounds connected to multiple lipid-related genes. *Solyc02g071700, Solyc02g090930, Solyc02g077430, Solyc06g053900, and Solyc08g063090* were among the genes with numerous connections to lipids. Genetic mapping and correlation-based network analysis revealed predominantly inter-class regulation, with loci controlling several classes of lipids (Fig. 3-5, 7, 8). The network reveals an intra-class regulation exemplified by the correlation between 16 glycerolipids and *Solyc09g009570* encoding a hexadecanal dehydrogenase.

This study combined lipidomics with mGWAS and family-based QTL mapping to identify novel lipid-metabolism genes and expand our understanding of genome-level regulation of lipid biosynthesis in tomato, establishing a direct genetic link between lipid metabolism and FA-VOC production in tomato fruits. Hence, those genes are likely to be useful in breeding for the improvement of palatability and nutrient contents.

## MATERIALS AND METHODS

### Plant material

The GWAS mapping population used in this study comprised a collection of 550 *S. lycopersicum* accessions that were selected after a phenotype-guided screen of over 7900 tomato accessions from around the world (Zemach et al., 2023). The GWAS panel contained modern cultivars, heirloom strains, and wild tomato species harvested from two independent greenhouse experiments during fall 2014 and fall 2015. The fall 2015 experiment (season) comprised a collection containing 388 *S. lycopersicum* accessions (cultivated tomato), 61 accessions of the *S. lycopersicum* var *cerasiforme,* 30 *S. pimpinellifolium* accessions, and 25 wild accessions. The 2014 season comprised a panel of 295 accessions including 24 *S. lycopersicum* var *cerasiforme* cherry tomato accessions and 271 cultivated varieties.

The *S. neorickii* BILs resulted from a cross between the green-fruited, self-compatible wild accession LA2133 and the processing tomato inbred variety cv. TA209 (*S. lycopersicum*). F_1_ hybrids were then backcrossed for three generations to cv. TA209 as described by Brog et al. (2019) followed by 10 generations of self-pollination to achieve BILs with maximum homozygosity of the wild genomic introgressions (Supplemental Fig. S11). The *S. neorickii* BIL population contained 142 lines. Due to poor germination, only 107 lines were used for this study. Additionally, hybrids for all BILs were produced in the background of the cv. TA209 recurrent parent to evaluate the wild introgressions in a heterozygous state.

For both the GWAS panel and the *S. neorickii* BILs population, pericarp tissue was isolated from ripe fruits, snap frozen in liquid nitrogen and stored at −80°C before extraction.

*Sl-LIP8*, *CFAPS1* KO lines and wild type Fla.8059 were grown in randomized, replicated plots in a heated greenhouse on the University of Florida campus or a field in Live Oak, Florida, using recommended commercial practices. All fruits for lipid and FA-VOC quantification were harvested at a full-red ripe stage.

### Lipid extraction protocol

Lipids were extracted from the GWAS panel harvested in two consecutive years from plants grown in the greenhouse and 2–4 independent biological replicates of *S. neorickii* BILs from fruit pericarp (Hummel et al., 2011). Briefly, 120 mg of ripe frozen fruits were used to make aliquots. Lipids were extracted with 1 ml of pre-cooled (– 20°C) extraction buffer (homogenous methanol:methyl-*tert*-butyl-ether (1:3) mixture + internal standards). After 10 min incubation in 4° C and sonication for 10 min in a sonic bath, 500 µL of water/methanol mixture was added. Samples were then centrifuged (5 min, 14000 x g). The lipophilic phase was collected and dried under vacuum.

### UPLC–FT–MS measurement

Samples were processed using ultra-performance liquid chromatography coupled with Fourier transform mass spectrometry (UPLC–FT–MS) on a C_8_ reverse phase column (100 x 2.1 mm x 1.7 µm particle size, Waters) at 60 °C. Samples were first subjected to liquid chromatography to separate them to their components. The mobile phase consisted of 1% 1 M NH_4_OAc and 0.1% acetic acid in water (Buffer A) and acetonitrile/isopropanol (7:3, UPLC grade BioSolve) supplemented with 1 M NH_4_OAc, 0.1 % acetic acid (Buffer B). The dried lipid extracts were re-suspended in 500 µL buffer B. The following gradient profile was applied: 1 min 45% A, 3 min linear gradient from 45% A to 35% A, 8 min linear gradient from 25 to 11% A, 3 min linear gradient from 11 to 1% A. After washing the column for 3 min with 1% A the buffer was set back to 45% A, and the column was re-equilibrating for 4 min, leading to a total run time of 22 min. The flow rate of the mobile phase was 400 µL/min.

Mass spectra were acquired using an Exactive mass spectrometer (Thermo Fisher, http://www.thermofisher.com) equipped with an ESI interface. All the spectra were recorded using altering full scan and all-ion fragmentation scan mode, covering a mass range from 100–1500C*m/z* at a capillary voltage of 3.0 kV with a sheath gas flow value of 60 and an auxiliary gas flow of 35. The resolution was set to 10,000 with 10 scans per second, restricting the Orbitrap loading time to a maximum of 100Cms with a target value of 1E6 ions. The capillary temperature was set to 150 °C, while the drying gas in the heated electrospray source was set to 350 °C. The skimmer voltage was held at 25CV while the tube lens was set to a value of 130 V. The spectra were recorded from min 1 to min 20 of the UPLC gradient.

### Targeted lipid profiling by LC–MS and data acquisition

Processing of chromatograms, peak detection, and integration were performed using REFINER MS® 10.0 (GeneData, www.genedata.com). Workflow comprised peak detection, retention time alignment, chemical noise and isotopic peaks removal from the MS data. Obtained mass features characterized by specific peak ID, retention time, m/z values, and intensity were further processed using in-house R scripts (Team, 2000). Clusters with mean signal intensities lower than 40,000 were removed and only peaks present in at least 80% of the samples were kept for analysis. Every day more than 65 samples were processed using LC-MS including quality controls at the beginning, the middle, and at the end of each day run. Peak intensities were weight- and day-normalized and log_2_-transformed. After that, obtained molecular features were queried against a lipid database for annotation. The in-house database used includes 219 lipid species of the following classes: diacylglycerols (DAGs), digalactosyldiacylglycerols (DGDGs), monogalactosyldiacylglycerols (MGDGs), phosphatidylcholines (PCs), phosphatidylethanolamines (PEs), and triacylglycerols (TAGs) (Giavalisco et al., 2011). Their retention times and exact masses were compared against those of the reference compounds, allowing maximal deviations of 0.1 minutes and 10 ppm. Identified lipids were confirmed by manual verification of the chromatograms using Xcalibur (Version 3.0, Thermo-Fisher, Bremen, Germany).

### General statistical and multivariate analysis of lipidomic data

Principal component analysis plot (Fig. 1, 7) boxplots analysis (Fig. 1, 3-7, Supplemental Fig. S6), pleiotropic map (Fig. 2), volcano plot (Fig. 3), heatmaps (Fig. 4, 6, 7) for the lipidomic data were obtained using R software version 4.3.1.

Complete pairwise correlations were calculated for lipid quantification from the GWAS panel of two consecutive years and gene expression information (Zhu et al., 2018). Correlations exceeding a significance threshold of 0.05 (adjusted by FDR) and Pearson correlation coefficient of ≥ 0.3 or ≤ 0.3, were assigned as edges in the network. Calculations were performed in R using package “Hmisc”, processed for visualization with the package “Rcy3” and visualized in Cytoscape 3.9.1 (see https://cytoscape.org/).

The chromosomal distribution of mQTL identified (Supplemental Fig. S5) was obtained by applying the RIdeogram R package to visualize and map genome-wide data on idiograms using R software version 4.2.2 (Hao et al., 2020).

Heatmaps (Supplemental Fig. S2-4) were created using Multi Experimental Viewer (MeV) software. Lipids were clustered using complete-linkage clustering. To detect the fold change in lipid-feature levels across all *S. neorickii* BIL lines and for the GWAS panel for each lipid feature, fold change was calculated by dividing the raw value of the peak height of a certain lipid by the average of values across all the lines for each trait.

### GWAS and linkage QTL mapping

Accessions for mGWAS were genotyped by sequencing (GBS) at Cornell University following an established protocol (Elshire et al., 2011) which yielded 16,526 SNP markers (Zemach et al., 2023) all marker data are deposited in the Phenome Networks database (http://unity.phenome-networks.com). The final SNP matrix (16,526) used for the analysis was obtained by filtering for minor allele frequency ≥ 5% (MAF ≥ 5%). Three principal components were included in the MLM to account for population structure and the SNP fraction considered for PCA. The kinship matrix and other parameters were set to default values. Association analysis was conducted using both 16,526 SNPs obtained from the GBS on 550 accessions and 1.8 million SNPs across 367 overlapping (same) accessions were previously characterized (Tieman et al., 2017). GWAS was performed using a compressed mixed linear model (MLM) (Z. Zhang et al., 2010) implemented in the Genome Association and Prediction Integrated Tool (GAPIT) in R (Lipka et al., 2012).

The *S. neorickii* BILs were genotyped with a 10K SolCAP single nucleotide polymorphism chip, and 3,111 polymorphic markers were used for mapping recombination breakpoints on the background of the physical map of *S. lycopersicum*. On average the BILs harbor 4.3 introgressions per line, with a mean introgression length of 34.7 Mbp, allowing the division of the genome into 340 bins and enabling rapid trait mapping (Supplemental Fig. S11). Linkage QTL mapping was done by applying “qtl” package version 1.40-8 that follows Haley–Knott regression (Brog et al., 2019). Each *S. neorickii* BIL and the appropriate controls of LA2133, TA209, and their F_1_ hybrid were genotyped using an Illumina 10K SNP chip (www.illumina.com). The genetic map contains 3,111 genome-anchored SNP markers that were investigated to be polymorphic between *S. neorickii* and cv. TA209. The markers were divided into 340 bins with an average length of 2.07 Mbp per bin and composited from an average of 9.15 SNPs per bin (Brog et al., 2019).

### QTL identification and candidate gene selection

We extracted SNPs associated with lipid species with significant *p-values*. For the mGWAS with GBS SNPs (N=16,526) data, the *p*-value was calculated according to the formula < 1/N (N=16,526). For mGWAS with 1.8 million SNPs data, the *p*-value was calculated according to the formula < 1/N (N=1,800,000) (Wu et al., 2016).

Candidate loci in the GWAS were identified based on the logarithm (base 10) of odds (LOD) score. A LOD score ≥ 4.5 was chosen as significant for mGWAS using GBS SNP data. A LOD score ≥ 6.5 was chosen as significant for mGWAS using 1.8 million SNP data. All genes in a given QTL were taken as putative candidates. Candidate genes were selected for validation based on their sequence homology with Arabidopsis genes related to lipid metabolism, their tissue-specific expression, and functional annotation.

The lipid profile of each *S. neorickii* BIL was compared (i.e., analysis of variance [ANOVA], using permissive threshold, p < 0.05) to the lipid content of TA209. If it was significantly different from the TA209 genotype, the introgression was considered as harboring an mQTL. Causal genes responsible for mQTL were identified considering the margins of introgressed regions from *S. neorickii* delimited by the genetic markers used in this work. The upstream and downstream borders of each introgression were established to be halfway between the inclusive and exclusive wild-species SNP (Brog et al., 2019).

### Linkage disequilibrium and haplotype analysis

Haplotype analysis was performed by using available SNP data. Accessions belonging to the same haplotype were grouped and the haplotype median was obtained for each lipid feature. To cluster haplotypes, a combination of allele sharing distance and Ward’s minimum variance was used (X. Gao & Starmer, 2007). One-way ANOVA for multiple comparisons was performed across all haplotypes (p < 0.05).

### Cloning of lipase-like protein (*TomLLP*) and generation of transgenic plants

Gateway® Technology (Invitrogen) was used in this work for overexpression of the lipase-like protein *(Solyc03g119980*) under the control of the Cauliflower mosaic virus *35S* (*CaMV35S*) promotor. Transformation of *S. lycopersicum* was performed as described in previous studies (Earley et al., 2006). The pDONR221 vector (4761 bp) for *attB-attP* reaction with a selectable marker for kanamycin resistance was used. The pK7WG2 (11,159 bp) vector for overexpression contained the *Solyc03g119980* gene in the sense orientation under the control of the *CaMV35S* promoter.

### qPCR gene expression analysis

Three parental M82 and six independent transgenic lines (with overexpressed *Solyc03g119980*) were grown in a greenhouse at the Max Planck Institute of Molecular Plant Physiology (Golm-Potsdam, Germany; Supplemental Data Set S10). The plants were grown in pots with 20 cm diameter three liter pots. At least three biological replicates were grown for each genotype. At least six fruit samples were collected from each replicate for transgenic lines and ten samples for the parental M82 line. Fruit pericarp was frozen in liquid nitrogen before storage at –80 °C. Frozen fruit pericarp was ground and RNA was extracted using a Thermo Fisher Scientific kit. The quality of extracted RNA was identified by visualization of the RNA integrity using 1% agarose gel and using a spectrophotometer (NanoDrop; Thermo Scientific). The final RNA concentration was adjusted to be equal in all the samples and cDNA was synthesized for each sample using a TAKARA cDNA synthesis kit for real-time gene quantification. Each cDNA sample was diluted 1:10, by adding RNAse-free water, and stored at –80 °C. The primers for *Solyc03g119980* were designed based on the Primer3 tool. The housekeeping genes were chosen according to the relevant literature (Supplemental Data Set S15) (Expósito-Rodríguez et al., 2008). The amplification efficiencies are indicated for every primer pair. The same cDNA in four dilutions (1:10; 1:100; 1:1000; 1:10000) was amplified with every primer pair (qPCR). The calibration curve slope (*R*^2^) was used for the identification of primer efficiency according to the formula:

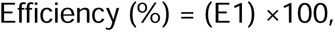

Where E was obtained from *R*^2^ regarding following formula:

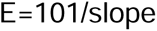

The differences in gene expression were calculated as fold change between independent transgenic lines and M82 according to the method of Schmittgen and Livak (Schmittgen & Livak, 2008) ΔCt value was calculated as the difference between the Ct (cycle threshold) of the candidate gene and the Ct of the control gene for normalization of gene expression, according to Schmittgen and Livak (Schmittgen & Livak, 2008).

### CRISPR lines

Three CRISPR lines for *TomLoxC (Solyc01g06540)* were created following the protocol described by Reem and Van Eck (Reem & Van Eck, 2019) with minor modifications. CRISPR constructs were created with a two-step golden gate cloning procedure to assemble a vector containing two guide-RNA-expressing cassettes, a kanamycin resistance gene, and the Cas9 nuclease, which was introduced into *Agrobacterium tumefaciens* GV2260. Guide RNAs were designed using the online tool CRISPR-P 2.0 (H. Liu et al., 2017). Transgenic plants were produced as previously described (Reem & Van Eck, 2019). Cas9-free plants homozygous for the gene of interest were transplanted in the greenhouse.

The design procedure and efficacy test of sgRNAs were performed using the CRISPR-P (http://cbi.hzau.edu.cn/cgi-bin/CRISPR) tool and the Guide-it^TM^ sgRNA *In vitro* Transcription and Screening System (Takara, Mountain View, CA, USA) according to the manufacturer. Vector construction and tomato transformation were performed as described previously (X. Li et al., 2020). Briefly, two 20-bp sgRNAs were inserted into a CRISPR/Cas9 binary vector (pCAMBIA2300_CR3-EF), in which the target sequence was driven by the Arabidopsis *U6-26* promoter and *Cas9* by 2 x 35S. The sgRNA sequences are listed in Supplemental Data Set S15. The final binary vector was transformed into cultivar Fla. 8059 by *Agrobacterium*-mediated transformation (X. Li et al., 2020). Genomic DNA was extracted from T1 and T3 homozygous *cfaps1* leaves and flanking regions containing the target sites were amplified using the specific primers *CFAPS1*-F and *CFAPS1*-R. The homozygous *cfaps1* allele was verified in the T1 and T3 generation by PCR-based sequencing. Cas9-free plants were used for quantitative analysis. The primers used for amplification and genotyping are listed in Supplemental Data Set S15.

Plant material from CRISPR lines targeting SI-LIP8 (*Solyc09g091050*) have been previously published (X. Li et al., 2020). The sgRNAs were inserted into the pCAMBIA2300_CR3-EF vector and transformed into Fla. 8059 by *Agrobacterium*-mediated transformation. Genomic DNA from an F2 plant backcrossed to WT and T3 tomato leaves was used for amplification with specific primers (Sl-LIP8-F and Sl-LIP8-R) for genotyping. Quantitative analysis used Cas9-free plants (X. Li et al., 2020).

### Experimental design for *TomLLP* (*Solyc03g119980*) validation

Sterilized seeds were grown on Murashige & Skoog selective plates with 20% sucrose (2MS) and Km under long-day conditions (16 h light, 8 h dark), temperature was kept at 21/16 °C (day and night, respectively), light intensity at 150 μE m^-2^ sec^-1^, humidity 75%. After 3 weeks, seedlings that survived selection were transported to the greenhouse in individual round pots with soil (potting compost) for fruit production and seeds for the next generation. The plants with empty-vector control and wild-type plants were grown in the same conditions as plants with overexpressed candidate genes.

### Experimental design of the Arabidopsis orthologue (*CSE, At1g52760*) of candidate gene (*TomLLP*, *Solyc03g119980*) validation

AtCSE_KD (*cse-1*, SALK_008202C) and AtCSE_KO1 (*cse-2*, SALK_023077) have been previously described (W. Gao et al., 2010; Vanholme et al., 2013) AtCSE_KO2 (GABI_368D11) is a T-DNA insertion mutant and was obtained from the GABI-Kat collection (Rosso et al., 2003). The T-DNA flanking sequence was analyzed via PCR with the primers 5’-ACCATTAGATGGTGAAATCAAAGG-3’ (1) and 5’-ATAATAACGCTGCGGACATCTACA-3’ (2), whereas the absence of the T-DNA was analyzed via PCR with the primers 1 and 5’-CTTGATAGCCTTCCCAACCA-3’ (3). The AtCSE_KO2 T-DNA insertion was confirmed to be positioned in the second exon (Vanholme et al., 2013).

---

**Supplemental Fig. S1.**
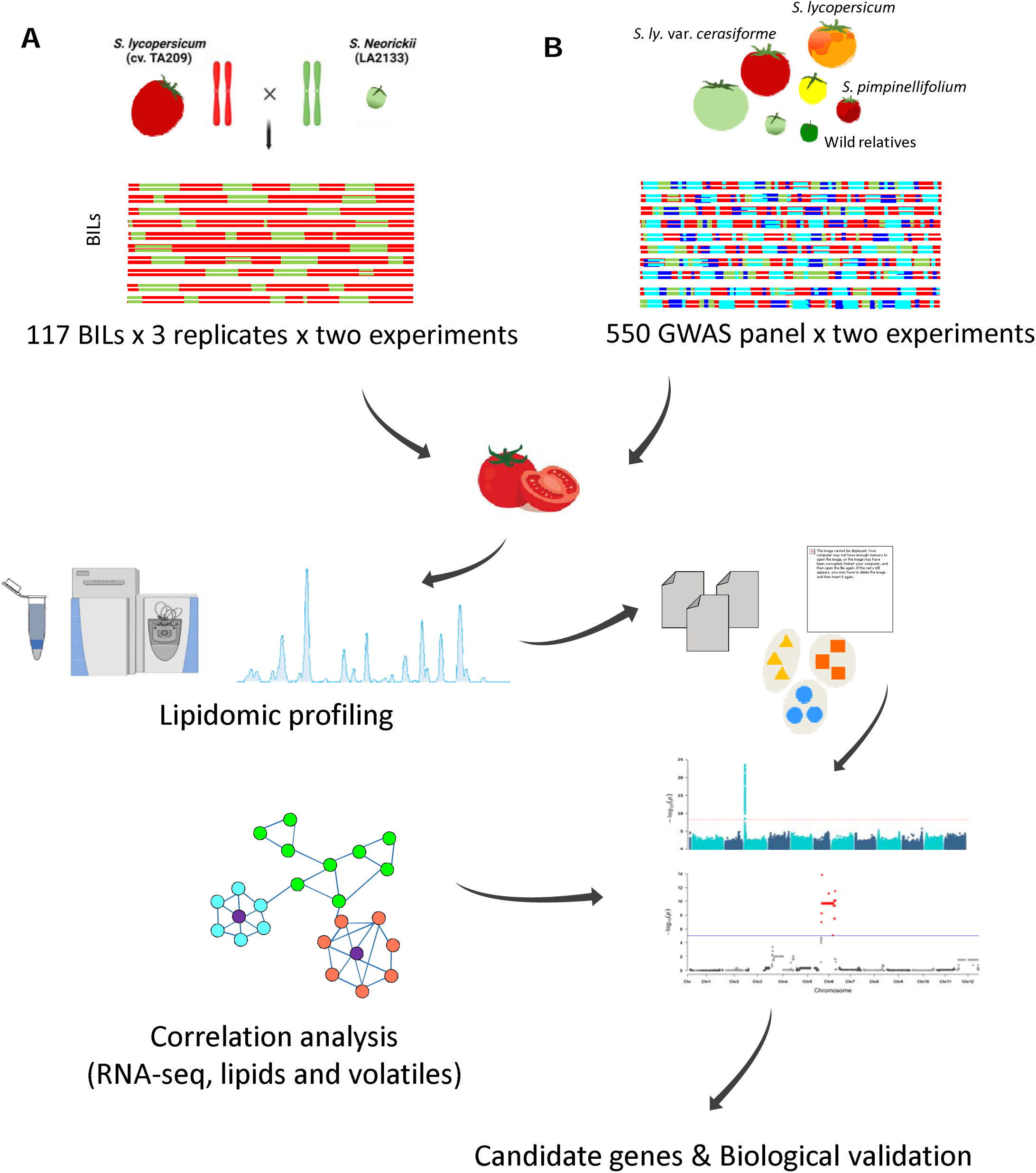
Schematic model of conducted experiments focused on investigation of genes underlying lipid metabolism in tomato fruit pericarp applying forward genetic approaches using association panels represented by **A)** *S. neorickii* biparental population, and **B)** unrelated cultivated tomato genotypes for genome-wide association study (GWAS).

**Supplemental Figure S2.**
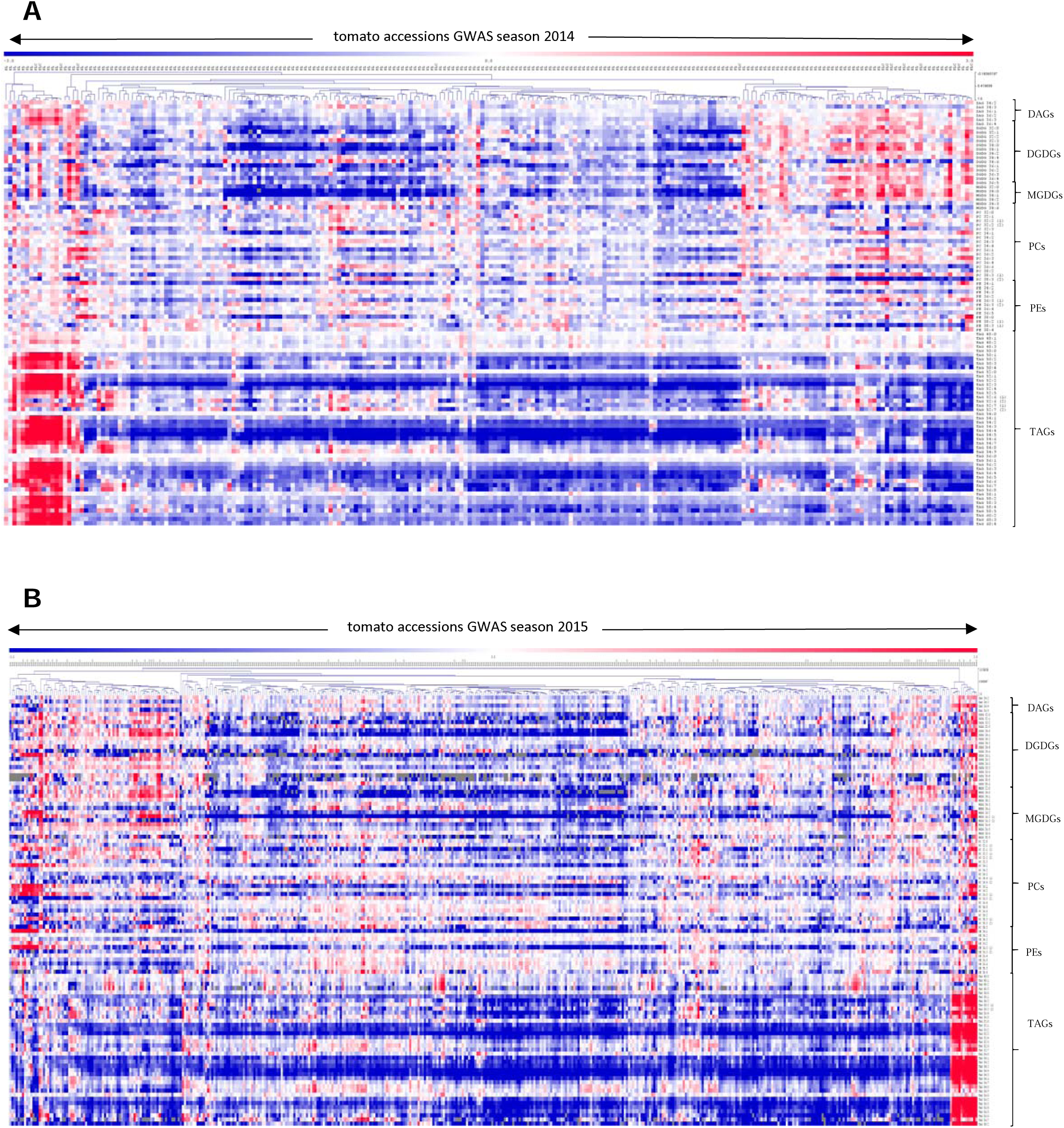
Heatmap of lipid levels across 550 accessions of the GWAS panel. The data represent lipidomic profiling of material harvested in two consecutive years 2014 **A)** and 2015 **B)** of plants grown in the greenhouse. For each lipid species mean lipid level was calculated and the level of the same lipid in each accession was normalized to this mean by dividing each lipid value by this mean. Each season was normalized separately and presented in a logarithmic scale (log2). Regions of red or blue indicate lower or higher compared to the average of each lipid species, respectively.

**Supplemental Figure S3.**
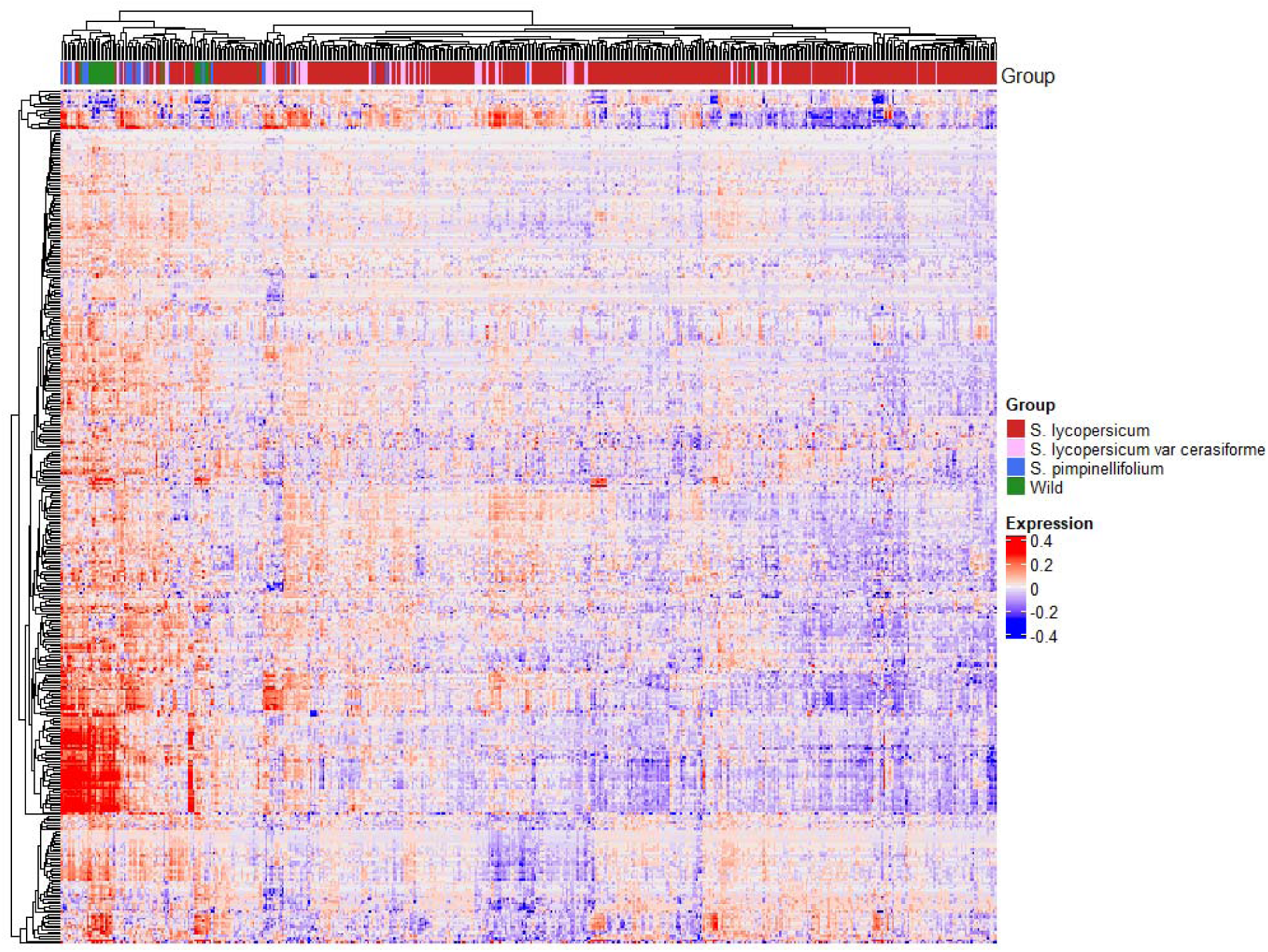
Heatmap of lipid levels across 550 accessions of the GWAS panel belonging to *S. lycopersicum*, *S. lycopersicum* var. *cerasiforme*, *S. pimpinellifolium*, and wild tomato groups. For each lipid species mean lipid level was calculated and the level of the same lipid in each accession was normalized to this mean by dividing each lipid value by this mean. The data are presented in logarithmic scale (log2). Regions of red or blue indicate lower or higher compared to the average of each lipid species, respectively.

**Supplemental Figure S4.**
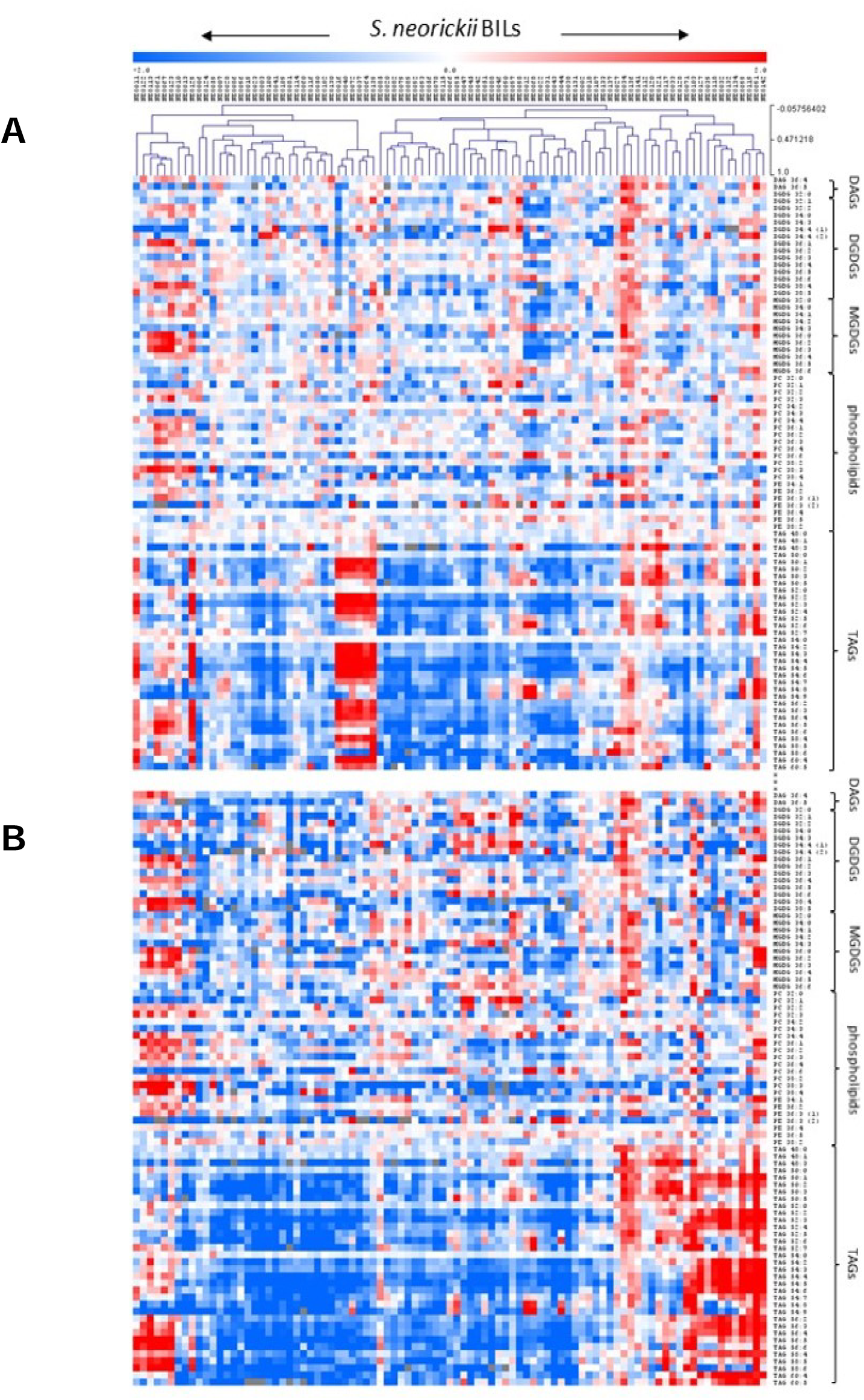
Heat map of lipid profiling across *S. neorickii* backcross inbred lines (BILs). The data represent lipidomic profiling of material harvested from *S. neorickii* BILs population **A)** heterozygous and **B)** homozygous lines. For each lipid species mean lipid level were calculated and the level of the same lipid in each BIL were normalized to this mean by dividing each lipid value by this mean. Each season was normalized separately and presented in a logarithmic scale (log2). Regions of red or blue indicate lower or higher compared to the average of each lipid species, respectively. Regions of white color, reflecting many of the chromosomal segment substitutions, do not affect lipid levels.

**Supplemental Figure S5.**
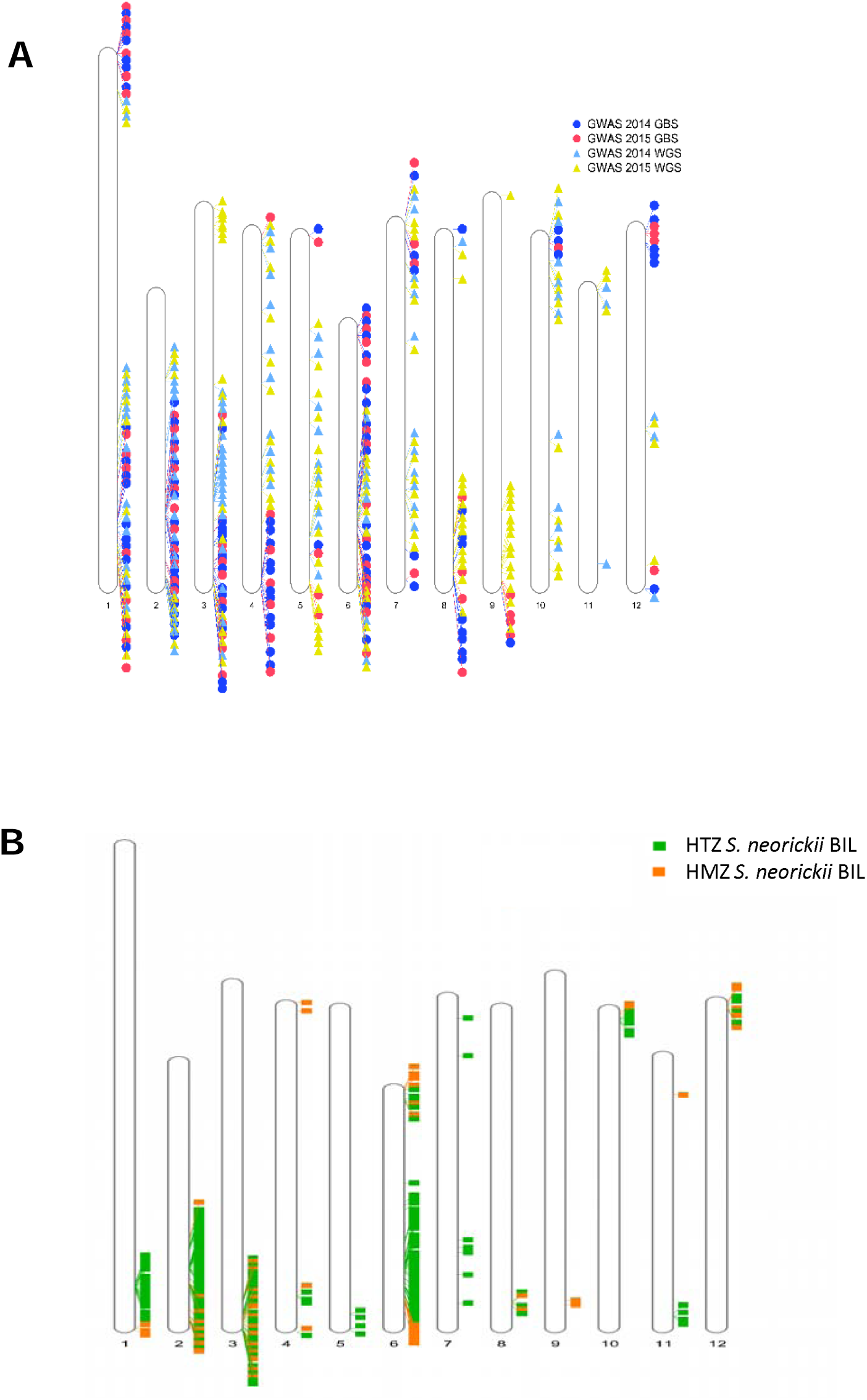
Chromosomal distribution of identified mQTL. **A)** Idiogram represents a chromosomal distribution of the mQTL resulting from GWAS of material harvested in two consecutive years using GBS and WGS SNPs data. **B)** Chromosomal distribution of the mQTL found in the BIL mapping of heterozygous and homozygous lines.

**Supplemental Figure S6.**
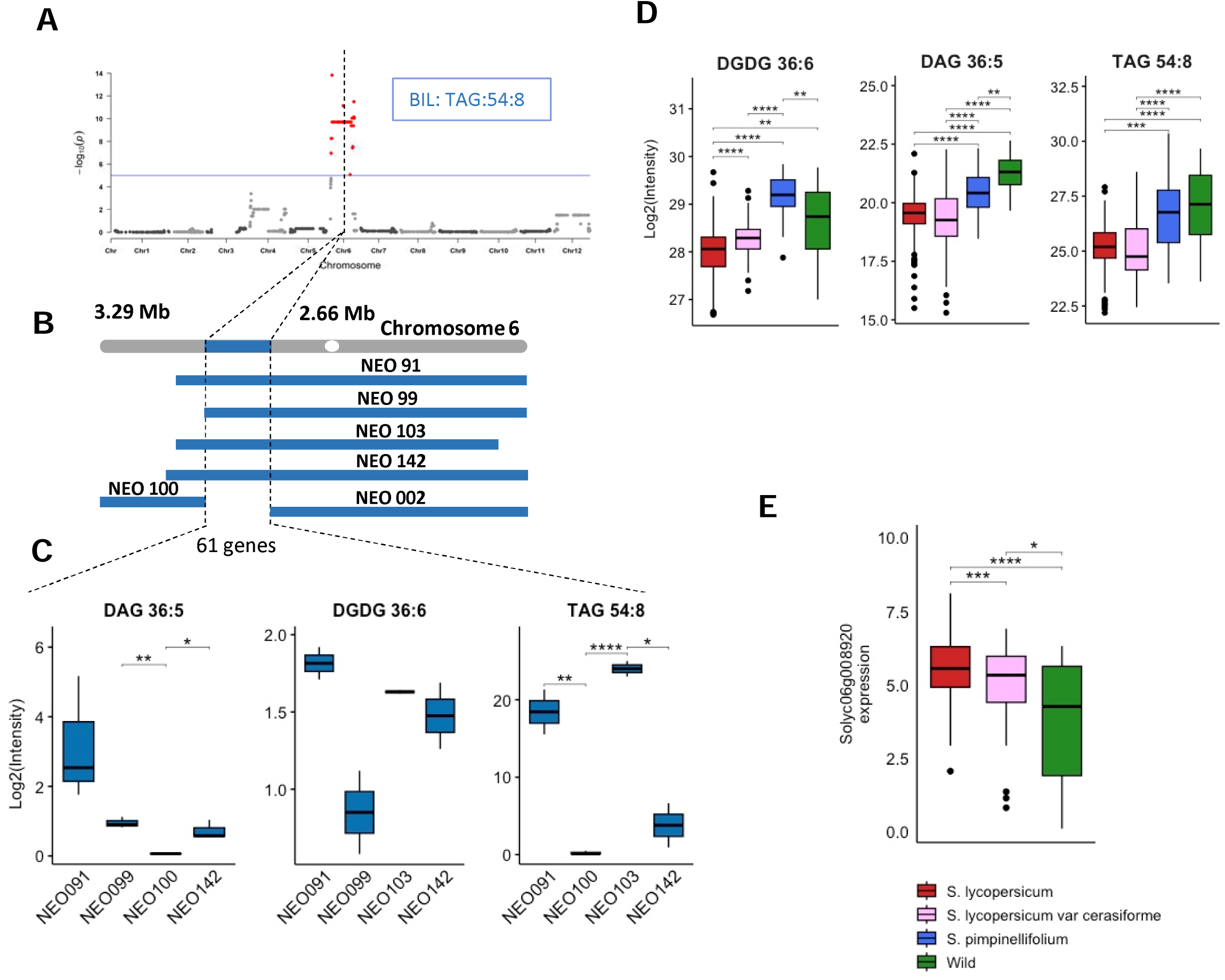
Linkage mapping of lipids in the BILs population reveled an mQTL comprising 61 genes harboring *Solyc06g008920,* encoding an Acetyl-CoA synthetase. **A)** Association plot of TAG 54:8 obtained with linkage mapping using *S. neorickii* BIL population. **B)** *S. neorickii* tomato segments introgressed into cultivated tomato variety TA209 on chromosome 6. **C)** Levels of DAG 36:5, DGDG 36:6, and TAG 54:8 in BILs sharing the *S. neorickii* introgression on chromosome 6 and BILs with the TA209 background. **D)** Average lipid levels for *S. lycopersicum* (n = 388)*, S. lycopersicum* var. *cerasiforme* (n = 61), *S. pimpinellifolium* (n = 30), and diverse wild tomato species (n = 25). **E)** *Solyc06g08920* transcript level in fruits of *S. lycopersicum* (n = 258)*, S. lycopersicum* var. *cerasiforme* (n = 56), and *S. pimpinellifolium* (n = 6). Significances (*p-value*) are indicated by letters using Student‘s t-test.

**Supplemental Figure S7.**
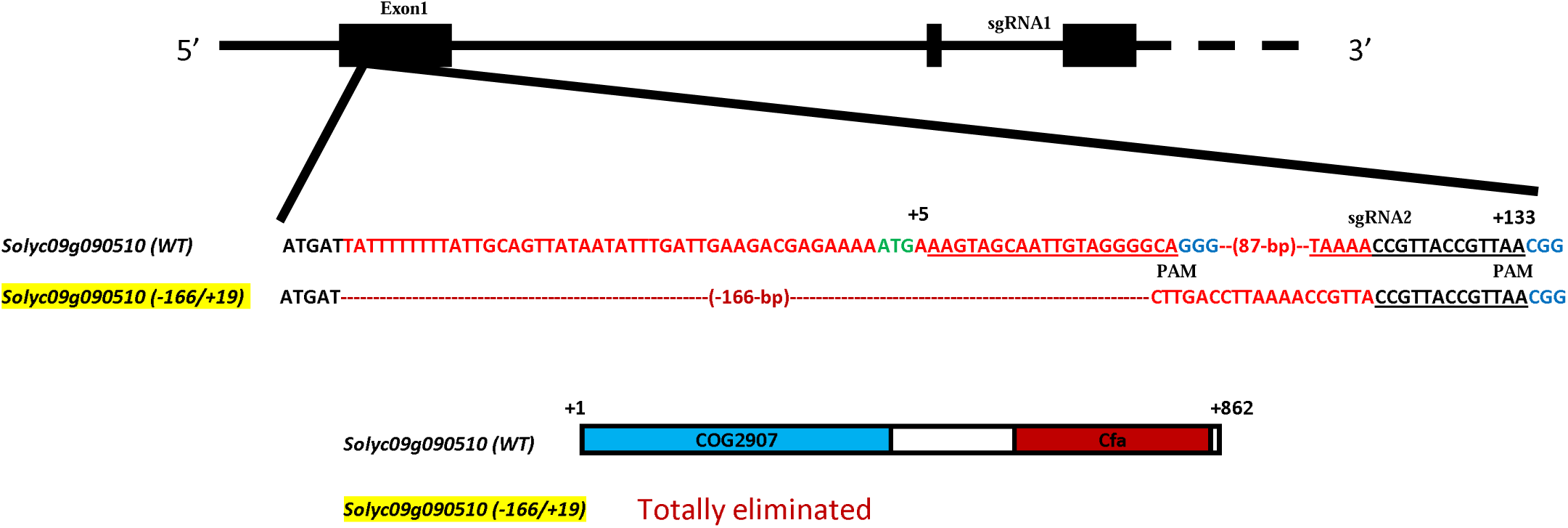
Construction and characterization of *CFAPS1*-edited lines. *CFAPS1* KO line (Fla. 8059 background) exhibits a deletion of 166 bp and an insertion of 19 bp in the first exon.

**Supplemental Figure S8.**
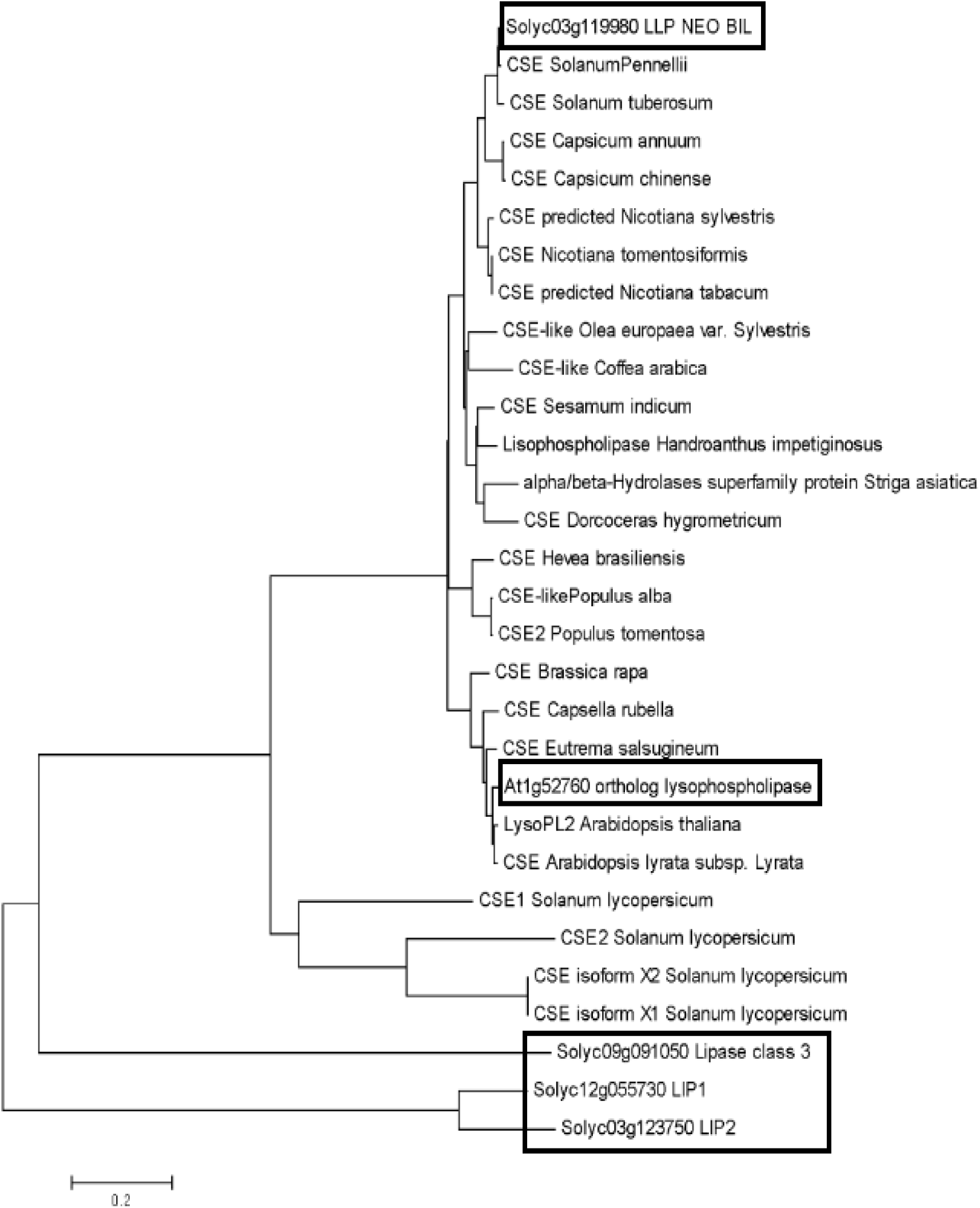
Phylogenetic analysis of the caffeoyl shikimate esterase (CSE) family. Coding sequences of genes with confirmed function as CSE or putatively annotated as CSE were used for the construction of a phylogenetic tree. Frame highlights two genes, the tomato *TomLLP* and *CSE* from Arabidopsis. Genes IDs are specified in Supplemental Data Set S9. A phylogenetic tree was reconstructed with the neighbor-joining method.

**Supplemental Figure S9.**
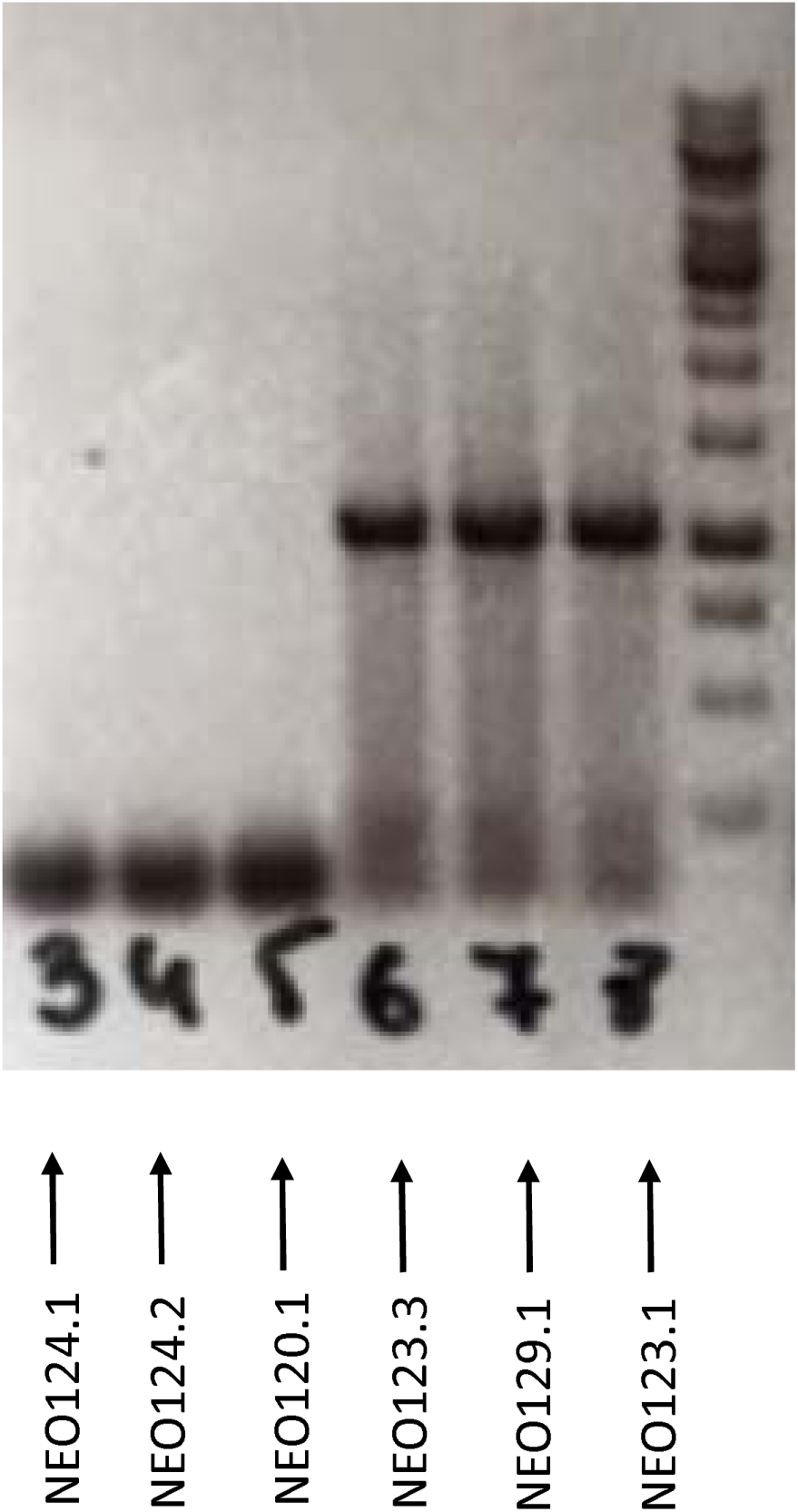
Expression level of *TomLLP* across six *S. neorickii* BILs. Monitoring the expression level of *TomLLP* using PCR on six BILs. Of which, in the region containing *TomLLP:* three BILs with the *S. neorickii* background and three BILs with the TA209 background.

**Supplemental Figure S10.**
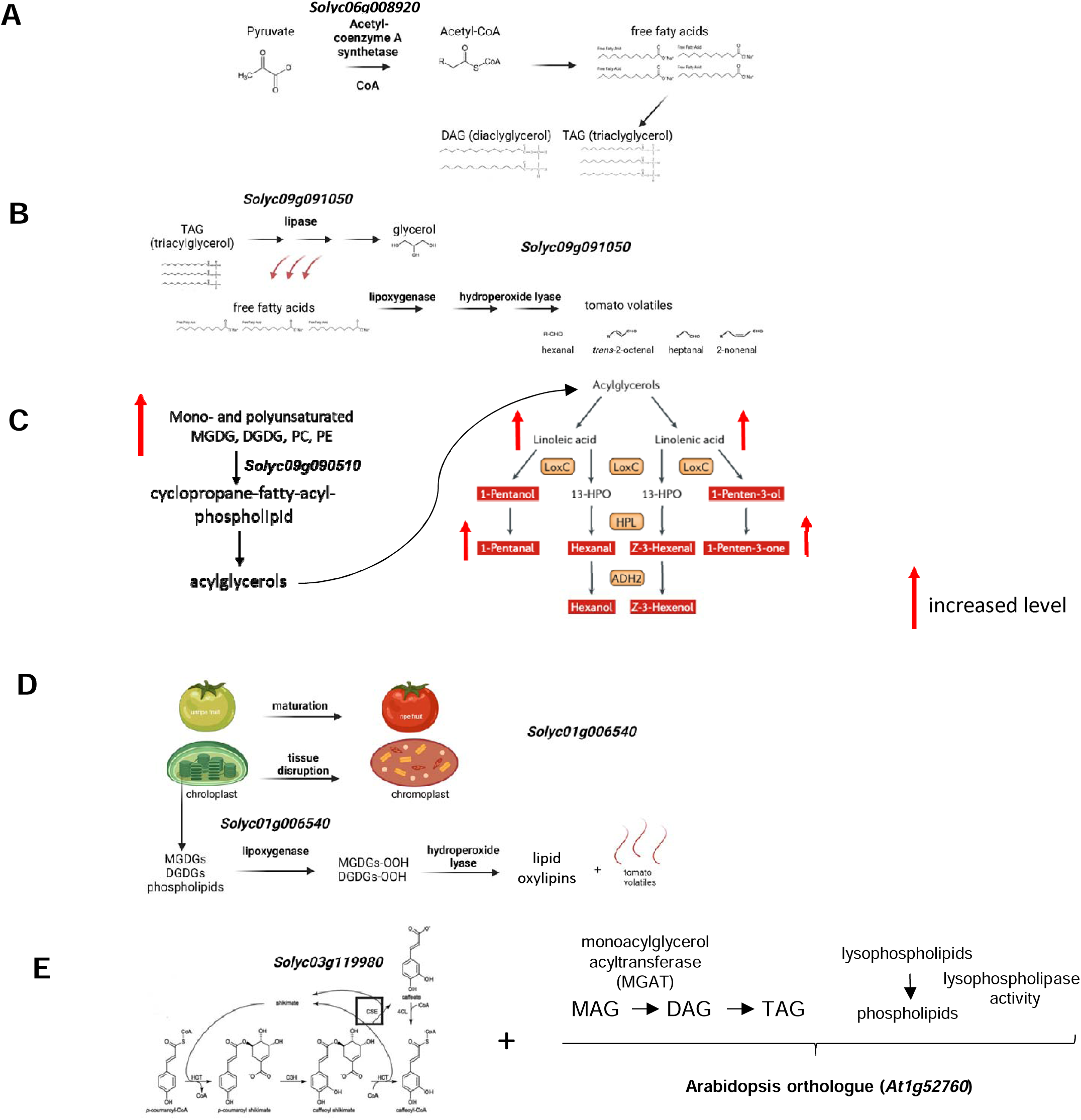
The metabolic pathways involve the identified lipid-related gene candidates. **A)** Schematic representation of the process of fatty acids synthesis by acetyl-CoA synthetase *(Solyc06g008920*) using pyruvate as a substrate. **B)** Schematic representation of the pathway of volatile synthesis from the free fatty acids liberated from triacylglycerol by class III lipase *(Solyc09g091050)*. **C)** Schematic representation of the process of conversion of membrane lipids (phospho- and galactolipids) to acylglycerols via cyclopropane-fatty-acyl-phospholipid with subsequent volatile production. **D)** Schematic representation of the process of lipid oxylipins and volatile production through the lipoxygenase enzymes *(Solyc01g06540)* in the lipase-independent pathway. **E)** The role of CSE (*At1g52760,* orthologue of *Solyc03g119980*) in the lignin biosynthetic pathway with additional identified acyltransferase and hydrolase activities.

**Supplemental Figure S11.**
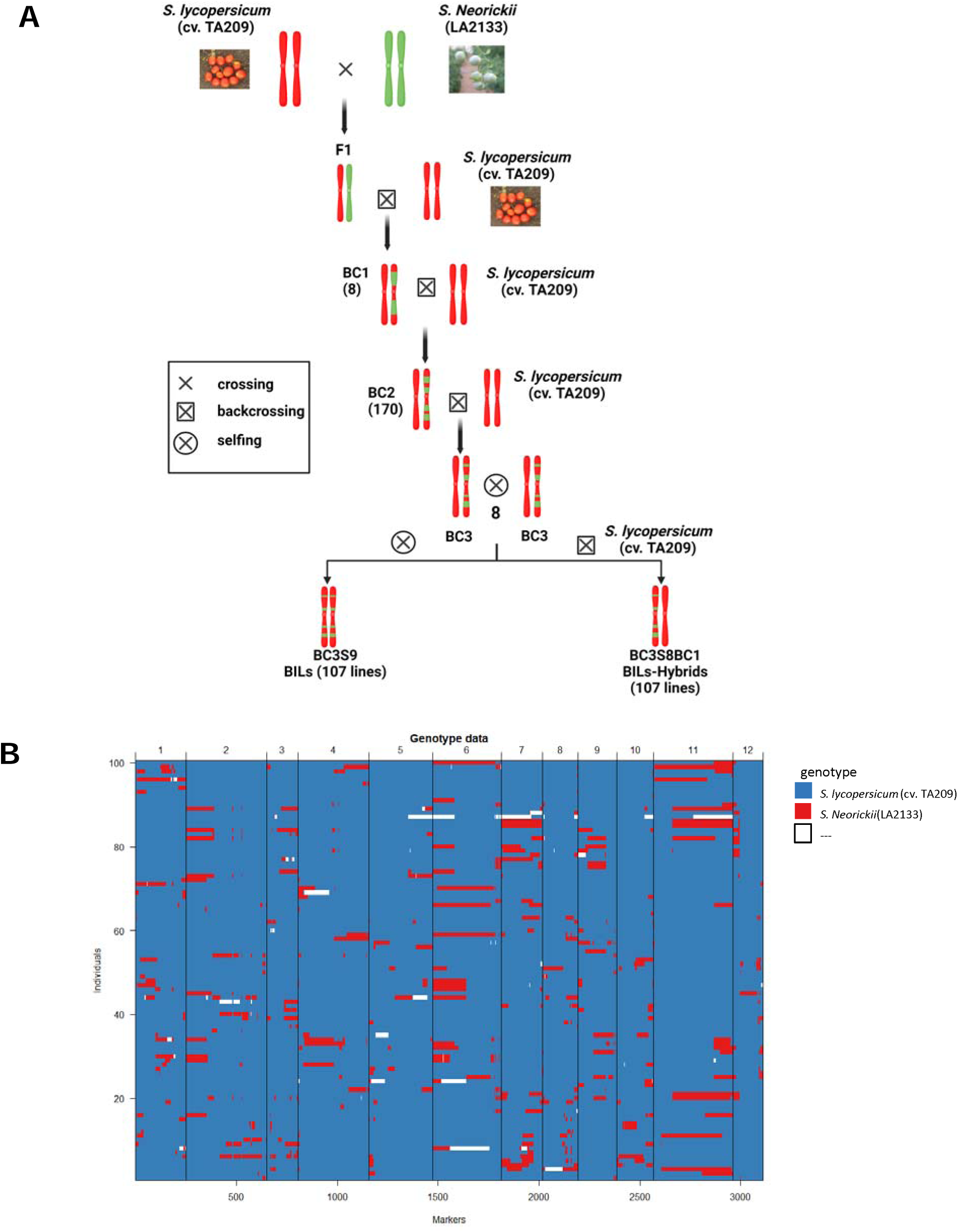
Breeding scheme and genetic map for backcross inbred lines (BILs). **A)** First cross: pollen from *S. neorickii* was placed onto the stigma of cv. TA209 to obtain F1 plants. Additional backcrosses with cv. TA209 was performed to decrease the *S. neorickii* genome introgression in the BILs. For each generation, the amount of plants is shown in parentheses. A final cross of the homozygous BILs with cv. TA209 was performed to obtain heterozygous lines (Brog et al., 2019). **B)** Schematic representation of *S. neorickii* backcrossed inbred lines. The BILs harbor on average 4.3 introgressions per line, with a mean introgression length of 34.7 Mbp, allowing the division of the genome into 340 bins and enabling rapid trait mapping (Brog et al., 2019).

## Supplemental Data Sets

**Supplemental Data Set S1**: Lipid content of the GWAS panel from 2015 harvest season.

**Supplemental Data Set S2**: Lipid content of the GWAS panel from 2014 harvest season.

**Supplemental Data Set S3**: Lipid content of homozygous *S. neorickii* BILs.

**Supplemental Data Set S4**: Lipid content of heterozygous *S. neorickii* BILs.

**Supplemental Data Set S5:** List of lipid compounds identified and quantified in both GWAS and *S. neorickii* BILs.

**Supplemental Data Set S6:** Number and distribution of the identified mQTL in both GWAS and *S. neorickii* BILs.

**Supplemental Data Set S7:** Lipids content in the *Sl-LIP8* KO and the wild-type tomato (c.v Fla.8059). ±SE; n = 4; Significances are indicated by * < 0.05, ** < 0.01, *** < 0.001 using Student’s t-test.

**Supplemental Data Set S8:** Lipid and FA-VOCs contents in the CFAPS1 KO lines and the control WT tomato. ±SE; n ≥ 10; Significances are indicated by * < 0.05, ** < 0.01, *** < 0.001 using Student’s t-test.

**Supplemental Data Set S9:** Coding sequences of CSE genes used for the construction of a phylogenetic tree.

**Supplemental Data Set S10:** Expression levels of Solyc03g119980 in OE lines using qPCR.

**Supplemental Data Set S11:** Lipids content in the *TomLLP* OE lines and the control WT (M82) tomato. ±SE; n = 5; Significances are indicated by * < 0.05, ** < 0.01, *** < 0.001 using Student’s t-test.

**Supplemental Data Set S12:** Lipids content in CSE knock-down, CSE knock-out lines *A. thaliana* wild type (A.th_WT). ±SE; n ≥ 4; Significances are indicated by * < 0.05, ** < 0.01, *** < 0.001 using Student’s t-test.

**Supplemental Data Set S13:** Lipids content in the TomLoxC KO lines and the wild-type tomato. ±SE; n=7; Significances are indicated by * < 0.05, ** < 0.01, *** < 0.001 using Student’s t-test.

**Supplemental Data Set S14:** Summary data sets of correlation-based network analysis

**Supplemental Data Set S15:** Primers used in this study

## Acknowledgments

S.A. acknowledges funding by the PlantaSYST project by the European Union’s Horizon 2020 Research and Innovation Programme (SGA-CSA nos 664621 and 739582 under FPA no. 664620), and NatGenCrop project: HORIZON-WIDERA-2022-TALENTS-01, No. 101087091. W.B. acknowledges financial support from the ERC Advanced grant POPMET.

## Authors’ contribution

A.K., X.L. performed experiments. A.K., X.L., J.L., JvS, M.B., E.K., K.K., L.R., M.W.A., K.G., A.K., A.-K. R. performed data analysis. A.D., A.I., provided technical and computer support. I.G., D.T., J.F., R.V., W.B., D.Z. provided plant materials. H.K., S.A. conceptualized the experiment. A.K., X.L., H.K., S.A. wrote the manuscript with input from all authors.

